# Detection of human unannotated microproteins by mass spectrometry-based proteomics: a community assessment

**DOI:** 10.1101/2025.02.19.639069

**Authors:** Aaron Wacholder, Eric W. Deutsch, Leron W. Kok, Jip T. van Dinter, Jiwon Lee, James C. Wright, Sebastien Leblanc, Ayodya H Jayatissa, Kevin Jiang, Ihor Arefiev, Kevin Cao, Francis Bourassa, Felix-Antoine Trifiro, Michal Bassani-Sternberg, Pavel V. Baranov, Annelies Bogaert, Sonia Chothani, Ivo Fierro-Monti, Daria Fijalkowska, Kris Gevaert, Norbert Hubner, Jonathan M. Mudge, Jorge Ruiz-Orera, Jana Schulz, Juan Antonio Vizcaíno, John R Prensner, Marie A. Brunet, Thomas F. Martinez, Sarah A. Slavoff, Xavier Roucou, Jyoti S. Choudhary, Sebastiaan van Heesch, Robert L. Moritz, Anne-Ruxandra Carvunis

## Abstract

Thousands of short open reading frames (sORFs) are translated outside of annotated coding sequences. Recent studies have pioneered searching for sORF-encoded microproteins in mass spectrometry (MS)- based proteomics and peptidomics datasets. Here, we assessed literature-reported MS-based identifications of unannotated human proteins. We find that studies vary by three orders of magnitude in the number of unannotated proteins they report. Of nearly 10,000 reported sORF-encoded peptides, 96% were unique to a single study, and 12% mapped to annotated proteins or proteoforms. Manual curation of a benchmark dataset of 406 manually evaluated spectra from 204 sORF-encoded proteins revealed large variation in peptide-spectrum match (PSM) quality between studies, with immunopeptidomics studies generally reporting higher quality PSMs than conventional enzymatic digests of whole cell lysates. We estimate that 65% of predicted sORF-encoded protein detections in immunopeptidomics studies were supported by high-quality PSMs versus 7.8% in non-immunopeptidomics datasets. Our work stresses the need for standardized protocols and analysis workflows to guide future advancements in microprotein detection by MS towards uncovering how many human microproteins exist.

## Introduction

Ribosome profiling (Ribo-Seq) studies have demonstrated widespread translation of short open reading frames (sORFs) outside of annotated coding sequences in eukaryotic genomes^1,2^, suggesting that the proteome may be much larger than currently annotated in databases such as UniProtKB.^3–6^ Several such individual sORF-encoded microproteins were experimentally found to be implicated in diverse biological processes across the tree of life such as muscle physiology and cancer.^7–12^ Yet, these well-characterized cases represent only a small fraction of the microproteins that could be encoded by translated sORFs.^13^ The translation products of many sORFs may be poorly conserved, of low abundance, or rapidly degraded, leading to uncertainty about their biological significance.^5,14,15^ There is a need, therefore, to identify the sORF-encoded microproteins that exist in the cell and have the potential to perform biological activities.

One systematic approach to identify unannotated microproteins predicted by Ribo-Seq is to search for peptide-level evidence in mass spectrometry (MS)-based proteomics or peptidomics datasets.^16,17^ In the typical case, a sequence database is constructed that consists of a curated protein sequence database (e.g. the UniProtKB human reference proteome^18^) joined together with a list of putative unannotated proteins (e.g. predicted products of translated sORFs cataloged by Ribo-Seq). This protein sequence database may then be used for analyzing conventional “shotgun” MS proteomics datasets, in which protein samples are digested using a protease, or for analyzing datasets generated by immunopeptidomics experiments, which attempt to identify peptides presented by human leukocyte antigens (HLAs) without requiring protease pretreatment.^19^ In both conventional proteomics experiments and immunopeptidomics experiments, the collected spectra will be generated from peptides derived from both annotated and unannotated proteins in the sample. Confident inference of an unannotated protein detection requires that the peptide uniquely supports an unannotated protein; i.e., that one can exclude the possibility that it derives from a protein in a curated protein sequence database. Detection confidence is generally controlled using a target-decoy approach^20^, which enables the calculation of a false discovery rate (FDR). The FDR can be set at the level of peptide-spectrum matches (PSMs), peptides, or proteins. Peptides and their inferred proteins passing the thresholds, usually 1% FDR at the peptide/protein level, are reported as detected.^21^ Protein-level MS evidence in a conventional proteomics experiment using trypsin or other proteases indicates that the protein existed in the cell. Immunopeptidomics can be used to validate Ribo-Seq predictions by confirming that an sORF was translated and the processed forms of its translation product was presented by HLA molecules, but cannot establish that the protein was stably present in the cell.^22^

Despite the promise of shotgun proteomics for rapid and large-scale microprotein identification, the small size, low abundance, atypical sequence characteristics and frequent transmembrane localization of microproteins pose major technical challenges for existing MS pipelines.^23–26^ For example, it can be impossible to observe multiple unique supporting peptides for microproteins whose sequence is too short to hold multiple cleavage sites, or if only one peptide is within the mass-over-charge range of the spectrometer. Therefore, the guidelines established by the Human Proteome Project^27^ for MS detection of proteins are difficult to apply fully, and researchers use a variety of ad hoc strategies.^16^ As the field develops and the number of reported microprotein detections grows, there is a need to assess which strategies are most effective for identifying genuine microproteins while minimizing false positives. Toward this goal, we brought together a group of experts to perform a systematic confidence assessment of previously reported unannotated protein MS detections.

## Results

### Reported numbers of unannotated proteins vary greatly between studies

To evaluate the extent to which unannotated proteins can be detected in proteomics data, our group of microprotein researchers assembled in 2023 to conduct a literature search for all papers reporting human unannotated protein detections published between 2019 and 2022. We identified 12 such studies published in this time window (Table 1). Seven studies searched for unannotated proteins in conventional proteomics data, while two studies searched for peptides derived from unannotated proteins in immunopeptidomics data, and three studies searched both classes of proteomics data. From each study, we obtained a list of the unannotated proteins reported to be detected (of any length), together with the PSMs supporting these detections (Supplementary Tables 1-2).

**Table 1:** Properties of reanalyzed studies. List of all studies reanalyzed. sORF database size indicates the number of sORFs in the protein sequence database in the MS analysis for each study. The number of these ORFs with proteomic support according to the study is also given. Considered noncanonical PSMs is the number of PSMs supporting a sORF-encoded protein reported in each study for which we could obtain the necessary information to evaluate; PSMs actually evaluated were selected randomly from this set. Annotation definition indicates the database used by each study to define the set of annotated or “canonical” proteins; all other proteins are considered to be unannotated, sORF-expressed proteins. Reported false discovery rate indicates the FDR given in each study for the list of sORF detections and whether this was calculated proteome-wide (a common FDR considering both unannotated and annotated proteins) or specific to the unannotated proteins.

| Citation | sORF database size | Considered noncanonical PSMs | Reported sORFs with MS support | HLA or non-HLA | Public datasets or new data generated | Source material | Annotation definition | Reported false discovery rate |
| --- | --- | --- | --- | --- | --- | --- | --- | --- |
| Cao et al. 2022<sup>30</sup> | Three-frame translation of transcriptome | 28 | 17 | non-HLA | New data | HEK293T | Human UniProtKB 2019 | 1% at peptide and protein level, proteome-wide |
| Bogaert et al. 2022<sup>28</sup> | 16,919 | 8 | 6 | non-HLA | New data | HEK293T cellular cytosol | Human UniProtKB/Swiss-Prot 2021 | <1% peptide, <2.5% protein, proteome-wide |
| Chothani et al. 2022<sup>4</sup> | 7,767 | 5,763 | 614 | non-HLA | Public datasets | NHDF and HUVEC (Slany et al. 2016<sup>31</sup>), ES (Shekari et al. 2017<sup>32</sup>), Heart (Doll et al. 2017<sup>33</sup>) | Human UniProtKB 2017 | 1% PSM level, unannotated specific |
| Duffy et al. 2022<sup>34</sup> | 38,187 | 2,445 | 366 | non-HLA | New data | Adult brain, Prenatal brain, hESC-derived neurons | Human UniProtKB | 1% at peptide and protein level, proteome-wide |
| Douka et al. 2021<sup>35</sup> | 45 | 18 | 8 | non-HLA | Public | SH-SY5Y cells (Murillo et al. 2018<sup>36</sup> and Brenig et al. 2020<sup>37</sup>) | Human UniProtKB 2019 | 10% at peptide level, proteome-wide |
| Prensner et al. 2021<sup>38</sup> | 553 | 6,236 | 140 | HLA and non-HLA | Public | 14 published mass spectrometry datasets | UCSC RefSeq | 1% at PSM level, proteome-wide |
| Ouspenskaia et al. 2021<sup>29</sup> | 237,437 | 9985 | 4903 | HLA and non-HLA* | Public and new | Lymphoblastoid cell line (Sarkizova et al. 2020<sup>39</sup>), patient-derived melanoma cell line, patient-derived glioblastoma cell line (Shraibman et al. 2019<sup>40</sup>), chronic lymphocytic leukemia tumor, ovarian carcinoma, renal cell carcinoma | Annotated genes on UCSC Genome Browser hg19 | 1% at PSM level, class-specific FDR for each type of unannotated ORF (e.g., uORF, dORF) |
| Chen et al. 2020<sup>41</sup> | 7,824 | 33 | 12 | HLA and non-HLA† | New data | iPSCs | Human UniProtKB | 1% at PSM level, proteome-wide |
| Chong et al. 2020<sup>42</sup> | Three-frame translation of transcriptome | 2,597 | 384 | HLA | New data | Patient-derived melanoma cell lines and lung cancer samples with matched normal tissues | Human UniProtKB/TrEMBL 2018 | Class-specific FDR for unannotated, keep only PSMs identified by both Comet and MaxQuant. Estimated FDR <0.001% |
| Martinez et al. 2020<sup>43</sup> | 7,554 | 1,160 | 319 | HLA | Public | Six cancer cell lines from Bassani-Sternberg 2015 (25576301): B-cells EBV transformed, B-cell leukemia, basal like breast cancer, colon carcinoma, primary fibroblast | Human UniProtKB/Swiss-Prot | 1% FDR at peptide level, proteome-wide |
| van Heesch et al. 2019<sup>6</sup> | 1,598 | 1,942 | 500 | non-HLA | Public and new | Heart (Doll et al., 29133944), iPSC-derived cardiomyocytes | Human UniProtKB 2017 | 1% targeted FDR, 50-60% estimated FDR |
| Lu et al. 2019<sup>44</sup> | 2,969 | 964 | 308 | non-HLA | New data | Cell lines: lung, colorectal cancer, liver cancer, cervical cancer | Human UniProtKB/Swiss-Prot | 1% FDR at PSM, peptide and protein level. |

\*Only HLA spectra were evaluated. †Only non-HLA spectra were evaluated.

A key motivation for initiating this community effort was the large variation in the number of validated unannotated proteins reported between studies, ranging from 6^28^ to 4,903^29^ (Figure 1A, Table 1). The peptides reported in support of unannotated proteins in each study were largely distinct: of 9,414 total reported peptides across the considered studies, only 326 (3.5%) were reported in more than one study. For 8 of 12 studies, fewer than 10% of the reported peptides were found in any of the other analyzed studies (Figure 1B, Supplementary Table 3). The low rate of replication is despite some studies analyzing the same collections of mass spectra, albeit with not fully overlapping databases of sORF sequences (Table 1). We do not interpret the high variability between studies as indicating that most reported detections are false: this high variability among reported detected peptides likely reflects in part the high variability in the size and composition of the sORF databases tested (Table 1)^16^ and the quantity of proteomic data analyzed, as well as the diversity of cell types examined, MS techniques used, HLA allotypes among the immunopeptidomics studies, and search algorithms. Nevertheless, in the absence of robust replicability to establish confidence, a closer assessment of the strength of evidence provided in each study for their reported detected unannotated proteins is needed.

**Figure 1:**
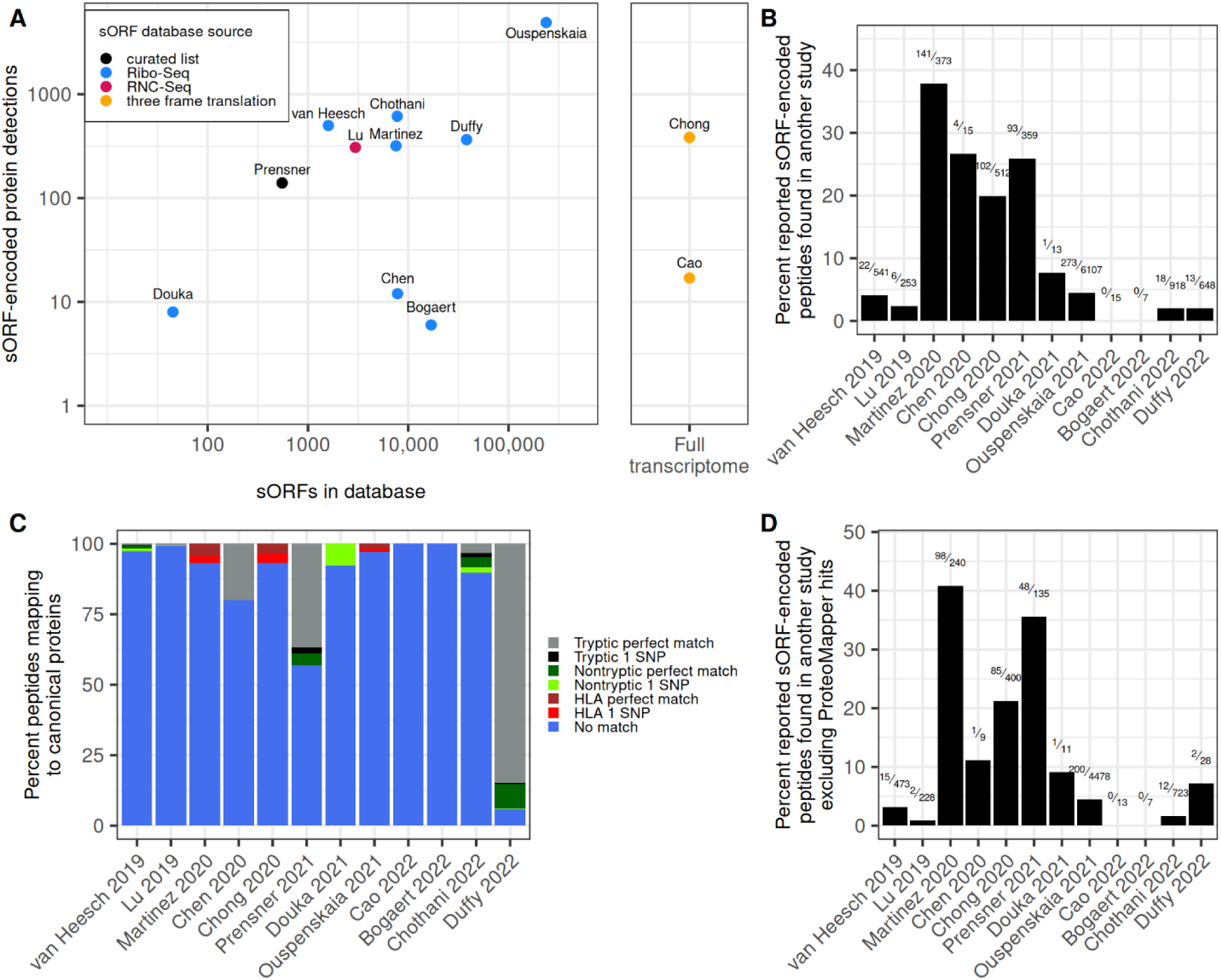
Broad variation among studies in reports of unannotated microprotein detection. A) The relation between the number of sORFs used to construct the protein database of each study and the number of sORF-encoded proteins reported detected by MS (Spearman correlation = 0.43, p = 0.2). Whether the sORF database was constructed using a curated list of known sORFs, all possible sORFs from three frame translation of a transcriptome, or a list of ORFs found to be translated using Ribo-Seq or RNC-seq data is indicated. B) For each study, the proportion of reported peptides supporting an unannotated protein that are also found by another study in our analysis is shown. The numbers of peptides found in other studies out of the total reported in the study are indicated above the bars. C) Proportion of peptides mapping to annotated proteins using the ProteoMapper tool, divided into categories depending on the number of common single nucleotide polymorphism (SNP) differences separating the peptide from the peptide present in the reference protein and whether the annotated peptide is tryptic; i.e., could be generated by cleavage after lysine or arginine. Semi-tryptic peptides (where only one peptide end is tryptic) are grouped with non-tryptic. Peptides from immunopeptidomics experiments were not generated by trypsin digestion and therefore are not classified as tryptic or non-tryptic. Peptides matching currently annotated proteins that were not annotated on UniProtKB/Swiss-Prot in 2016 (i.e., recently annotated proteins) are excluded. D) For each study, the proportion of reported peptides supporting an unannotated protein that are also found by another study in our analysis, excluding peptides that match to annotated proteins according to the ProteoMapper tool. Note that most studies have focused on different biological systems, which can limit the overlap.

### Do reported peptides uniquely support an unannotated protein?

We first assessed whether PSMs reported as evidence for the detection of an unannotated protein may also be attributed to an annotated protein. All the studies in our meta-analysis attempted to exclude potential annotated protein-matching peptides, but different analysis pipelines were implemented that might not have equally accounted for the full space of potential proteoforms of annotated proteins.^16^

To assess whether some peptides reported to derive from an unannotated protein could potentially be attributed to an annotated protein, we used the PeptideAtlas ProteoMapper^45^ tool. ProteoMapper takes neXtProt^46^ reported amino acid variants into account; i.e., it will find matches not just to the reference proteome but to proteins that differ from the reference by one or more variant amino acids. We restricted our analysis to peptides that differed from the reference sequence by at most one single amino acid variant. Given this restriction, 12% of peptides reported to support detection of an unannotated protein (1,161 of 9,732) also had a putative match to an annotated protein on ProteoMapper, with this rate varying from 0% to 96% across individual studies (Supplementary Table 1).

Recent updates in annotation could potentially explain why some reported peptides mapped to annotated proteins when we conducted this ProteoMapper search in 2023. To evaluate this possibility, we checked whether these annotated proteins were annotated in the 2016 version of UniProtKB/Swiss-Prot^18^, as all studies in our analysis used protein databases published after 2016 to define their annotated set (Table 1). Only eight distinct annotated proteins matching reported unannotated peptides in 2023 were absent from UniProtKB/Swiss-Prot in 2016, indicating that annotation updates are not a major explanation for peptides reported to support unannotated proteins mapping to annotated proteins.

Peptides reported to support unannotated proteins might also map to annotated proteins if the studies did not account for non-tryptic peptides or protein variants. We therefore divided the peptides mapping to annotated proteins by whether they were perfect matches to the UniProtKB/Swiss-Prot reference protein or differed by one single amino acid variant, and by whether they were predicted tryptic (i.e., peptides that could be generated by cleavage after arginine or lysine residues) or non-tryptic (including semi-tryptic) (Figure 1C). We note that some peptides in Chong et al. 2020^42^ map to both unannotated proteins and common variants of annotated proteins, but since this study used customized databases of annotated proteins reflecting each patients’ sequenced genotypes, these common variants were shown to be absent in the patient samples. Without such a customized database, it is difficult to fully rule out an annotated protein source given the possibility of unknown variants of annotated proteins, especially in cell lines or cancer samples.

For two studies, Prensner et al. 2021^38^ and Duffy et al. 2022^34^, a substantial fraction of reported unannotated peptides (10% or more) were perfect matches to tryptic peptides in reference proteins. The relatively high rate of matching UniProtKB protein references in Prensner et al. 2021^38^ might be explained by either the use of the UCSC RefSeq database to define the set of annotated proteins rather than UniProtKB, which was used by most other studies (Table 1), or not preferentially allocating all shared peptides to the annotated set. For Duffy et al. 2022^34^, spectra searches were conducted against custom databases of both annotated and unannotated proteins inferred to be expressed in the specific type of brain tissue or cell based on Ribo-Seq data, while all other studies included the full set of human annotated proteins in their protein database. Likely, annotated proteins not detected by Ribo-Seq may still be present in the sample, leading to peptides from annotated proteins potentially being falsely assigned to unannotated proteins. For two other studies^6,41^, more than half of reported peptides that mapped to both unannotated and annotated proteins were non-tryptic (Figure 1C). A peptide with a match to an annotated protein does not uniquely support an unannotated protein detection, even if the match is non-tryptic, as trypsin does not have perfect specificity and can vary in grade, cleavage could have been induced by other proteases (e.g. upon lysing cells and tissues), and protein processing in cells can yield non-tryptic peptides.

Overall, these results indicate a need to consider non-tryptic peptides and possible amino acid variants of annotated proteins to ensure that peptides uniquely map to an unannotated protein. Excluding potential hits to annotated proteins can be done with tools such as ProteoMapper^45^ or the neXtProt peptide uniqueness checker^47^, as suggested by the HUPO-HPP MS data interpretation guidelines^27^, or, ideally, using sample-specific customized protein sequence databases based on sequenced genotypes.

After excluding all reported peptides that mapped to annotated proteins according to ProteoMapper, the general trends we observed for the entire set of reported peptides supporting unannotated protein detections remained: for 8 of 12 studies, at least 90% of reported unannotated peptides were only reported in that study (Figure 1D). Therefore, we next examined the level of support PSMs provided for claimed unannotated protein detections.

### Assessing PSM quality by manual evaluation

To assess PSM quality among literature-reported peptides supporting detection of unannotated proteins, a random sample of PSMs from each study was manually evaluated by a panel of six expert evaluators. A total of 406 PSMs from 12 studies were evaluated (1.3% of total), corresponding to 307 peptides from 204 unannotated proteins. These PSMs were sampled after excluding peptides mapping to annotated proteins or proteoforms (Figure 1C). Of these 406 PSMs, 155 were evaluated by two evaluators each to enable determination of the overall consistency between evaluators. Additionally, a common set of 10 negative control PSMs was included in each sample, consisting of high-scoring decoy-spectrum matches intended to mimic PSMs that perform relatively well according to algorithms. Each PSM was rated on a scale of 1-5. Full evaluation criteria along with example spectra and explanations of their rating are given in Appendix 1. The PSMs assigned to each evaluator were ordered randomly and the evaluators were not informed as to the source publication of each PSM (Supplementary Table 4).

Agreement among evaluators was generally high. For the PSMs rated by two evaluators, ratings were well correlated (r = 0.82, p < 10^-10^) (Figure 2A). Only 14 of 155 (9%) PSM scores differed by more than one point. The negative controls scored consistently poorly (average score of 1.5), as expected. Evaluator ratings were also well correlated (r = 0.74, p < 10^-10^) with the dot product between the observed spectra and the spectra predicted by MS2PIP (Supplementary Figure 1).^48^ Among immunopeptidomics studies, PSMs with peptides that were predicted to bind to MHC molecules by NetMHC^49^ were rated more highly (n = 71, mean rating 3.94) than those with peptides not predicted to bind (n = 14, mean rating 3.29, p = 0.037 by two-sided permutation test, Supplementary Figure 2, Supplementary Table 5), consistent with manual evaluation discriminating between true and false discoveries. To investigate consistency between manual ratings and machine learning methods for spectral prediction, we generated predicted spectral libraries for all evaluated PSMs under several models using Oktoberfest (see Methods).^50^ We observed a moderate correlation between the best spectral angle between the model-predicted and experimental spectra (a measure of spectral similarity) and evaluator rating (r = −0.56, p < 10^-10^, n=274, Figure 2B), suggesting both similarities and differences in how expert evaluators and this spectral prediction method assess PSM quality.

**Figure 2:**
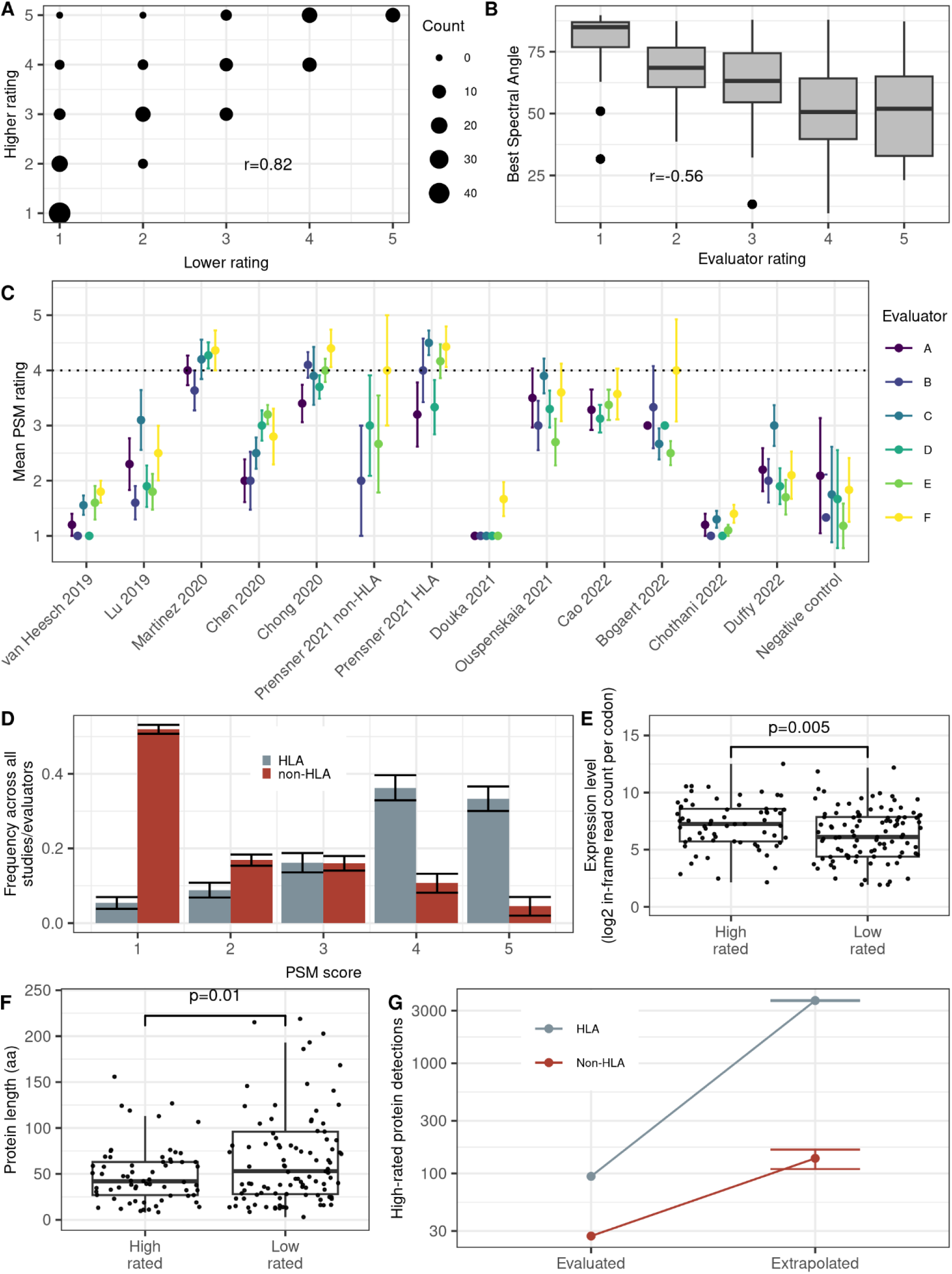
Expert manual evaluation of literature reported unannotated protein detections in mass spectrometry datasets. A) Counts of each pair of ratings among the PSMs that were assessed by two evaluators (n=155). The Pearson correlation between pairs of ratings is indicated. B) For each manually evaluated PSM, the spectrum was also predicted using several machine learning models (see Methods). The spectral angle is an indicator of how different the observed PSM was from the closest predicted spectrum, with larger angles indicating a worse match. The best spectral angles are indicated among PSMs grouped by evaluator rating. C) Mean ± standard error of ratings of PSMs sampled from each study, per each of six evaluators. Standard errors were corrected for finite population (total count of reported PSMs supporting unannotated proteins in the study). Ratings were given on a 1-5 scale. D) Overall distribution of ratings for unannotated protein PSMs among all studies and evaluators. Bars indicate standard errors. E) Log Ribo-Seq read counts for ORFs expressing proteins in PSMs rated highly (>3, n=65) or lowly (<3, n=105). Reads are from a collection of human Ribo-Seq studies (see Methods). F) Predicted lengths of proteins rated highly (>3,n=65) or lowly (<3, n=105). G) Evaluated and extrapolated counts of HLA and non-HLA high-rated (rating of 4 or 5) protein detections. Extrapolated counts give the number of high-rated protein detections expected if the entire dataset had been evaluated.

There was also a general consistency between evaluators in average rating per study (Figure 2C). The evaluated PSM quality varied across studies, with average rating ranging from 1.0 to 4.1 (Figure 2C). Three studies had average PSM ratings that did not exceed the negative controls. For one of these studies, van Heesch et al. 2019^6^, the authors recognized the high FDR in their search results, which led them to develop a customized strategy for estimating a microprotein-specific FDR and to favor selected reaction monitoring (SRM) for their downstream analyses. We did not evaluate these SRM results but focused solely on the reported shotgun proteomics hits. For Douka et al. 2021^35^, the low ratings are understandable because, rather than using a 1% FDR threshold, this study used a 10% threshold in anticipation of the low abundance of microproteins. For Chothani et al. 2022^4^, unannotated protein PSMs were identified by searching hundreds of MS runs individually with a 1% FDR threshold after removing all matches to the annotated proteome, then assembling the hits into a master list. A likely explanation is that, since spectra matching annotated proteins were removed prior to searching for unannotated proteins, there were few genuine detections in the MS runs analyzed. Under conditions of few genuine detections, it is difficult to precisely estimate FDR, leading to potential false positives (Supplementary Figure 3).^51^ Chothani et al. highlighted peptides found in multiple datasets; these peptides were not separately evaluated here.

The immunopeptidomics studies (Ouspenskaia et al. 2021^29^, Martinez et al. 2020^43^, and Chong et al. 2020^42^, and some peptides from Prensner et al. 2021^38^) reported substantially higher quality PSMs than most of the other studies (mean rating 3.8 vs. 2.3, n=13, p = 0.024 for difference in mean by two-sided permutation test, Figure 2C-D). The three studies that focused on HLA data have average scores above three, as do the HLA PSMs (but not non-HLA PSMs) from Prensner et al. 2021.^52^ The only non-HLA studies with average scores of three or more were Cao et al. 2022^30^ and Bogaert et al. 2022^28^, which reported only 28 and 8 PSMs derived from unannotated proteins, respectively (Figure 2C, Table 1). Overall, most (70%) evaluated PSMs supporting unannotated protein detections from HLA studies received a rating of at least 4, the threshold for convincing evidence of detection (See Appendix, Figure 2D). In contrast, only 15% of ratings for reported matches in non-HLA data were in the 4-5 range. These results are consistent with a recent study, Deutsch et al. 2024, where MS searches for peptide-level evidence supporting Ribo-Seq identified sORFs also found higher support in HLA than non-HLA datasets.^53^

Among 98 high-rated HLA peptides, 33 were reported in multiple studies, and 37 were validated by Deutsch et al. 2024 (1 supporting an ORF in Tier 1A, 26 in Tier 1B, and 10 in Tier 2B, Supplementary Figure 4). Of the 28 high-rated PSMs from non-HLA data, two involved peptides that were reported in multiple studies. Both peptides derive from the same sORF, located in the 5’ UTR of the MKKS locus. The protein encoded by this sORF (UniProt identifier Q9HB66 in UniProtKB/TrEMBL) has now accumulated enough peptide-level evidence to have become annotated as “core canonical” in PeptideAtlas in 2025, though it remains unannotated in UniProtKB/Swiss-Prot so far. Two high-rated non-HLA peptides were also identified as having strong evidence in Deutsch et al. 2024.^53^ These peptides mapped to the sORFs c11riboseqorf4 in the Tier 1A class (the highest level of support that an ORF is protein-coding) and c12norep33 in the Tier 2A class (weaker support). These observations illustrate how searching multiple sources of MS data contributes towards a more comprehensive view of sORF-expressed proteins and improves annotations of the human proteome.

### Higher rated PSMs are derived from more highly expressed sORFs

To assess whether our PSM ratings were influenced by the expression levels of the corresponding proteins, we compiled a large collection of human Ribo-Seq studies and analyzed translation levels harmoniously, using the iRibo program, for all the sORFs corresponding to evaluated PSMs for which genomic coordinates were provided by the original studies (191 sORFs; see Methods, Supplementary Tables 6-7).^54^ We found that reported unannotated proteins with corresponding PSMs rated 4 or 5 were more highly translated than those with corresponding PSMs rated 1 or 2 (difference in log Ribo-Seq read count per codon by two-sided permutation test, p = 0.005, Figure 2E). This is consistent with more highly expressed proteins being more readily detectable by MS and thus generating higher quality PSMs.^55^ Unexpectedly, high-rated proteins were also shorter on average by 37 amino acids than low-rated proteins (two-sided permutation test, p = 0.01, Figure 2F). There was no significant correlation between log iRibo p-value, indicating level of confidence that the ORF is translated, and PSM rating (r = 0.098, p = 0.18).

### Discovery of potential unannotated proteins

We next estimated the number of unannotated proteins we would expect to have strong MS support had we evaluated all reported detections. To do this, we extrapolated the number of unannotated protein detections that would be supported by high-scoring PSMs had we evaluated all PSMs among all studies, assuming the frequency of scores for each study would be the same as in the tested set (Figure 2G). Among unannotated proteins reported in non-HLA data, 27 evaluated proteins were supported by at least one PSM rated 4 or 5. We predict 137 of 1,749 (7.8%) would be supported by PSMs of this quality across the whole aggregated dataset. For HLA data, 94 evaluated proteins were supported by at least one PSM rated 4 or 5; we predict 3,706 of 5,705 (65%) would be found across the entire dataset. Other unannotated proteins are likely detectable in datasets outside our study scope. Thus, there is considerable potential for discovery even in the particularly challenging case of finding unannotated proteins in conventional enzymatically digested samples.

## Discussion

Given the growing recognition of the importance of microproteins in human health^56^, there is an urgent need to prioritize sORF-encoded microproteins that are supported by MS evidence. Here, we reanalyzed twelve published studies that reported detection of unannotated microproteins with MS. While most reported PSMs (70%) in immunopeptidomics studies were of high quality, around 85% of non-HLA PSMs were evaluated by a panel of proteomics experts to be of too low quality to provide evidence of peptide detection. These results point to a need for caution in interpreting claimed unannotated protein detections reported in the literature and motivate technological improvements for the evaluation of microprotein evidence moving forward. Many unannotated protein detections do appear strong, and the microprotein literature has provided great value in expanding the protein universe with real discoveries of likely biological significance.^53^ However, the idea that several hundreds to even thousands of unannotated proteins are genuinely detected in existing mass spectrometry datasets of conventional trypsin digests reflects an unrealistic expectation about the extent to which current shotgun proteomics can validate sORFs identified by Ribo-Seq. Why do immunopeptidomics studies identify many high-quality PSMs supporting unannotated protein detections while studies using conventional enzymatic digests identify only few? Many unannotated sequences found to be translated by Ribo-Seq lack signatures of evolutionary conservation and may not encode proteins that provide any benefit to the organism.^5,15,57^ It is plausible that many of these poorly conserved proteins are expressed but quickly degraded, and so can be found only as peptides bound to HLAs.^14,58^ However, there are also technical explanations for why HLA-bound peptides derived from unannotated microproteins may be easier to detect. Immunopeptidomics concentrates peptides bound to HLAs, which decreases sample complexity and may thereby enrich for low abundance microproteins. HLA peptides also have physico-chemical properties different from tryptic peptides that may affect detectability. Most immunopeptidomics datasets are from cancer samples, and some proteins may be expressed in some cancers but not in normal physiological conditions. Furthermore, microproteins may preferentially reside in cellular compartments that are hard to sample through non-HLA MS, such as membranes.^26^ Moreover, the laboratories that perform immunopeptidomics are often distinct from those that analyze non-HLA data and may differ in their sample preparation techniques, experimental setup, and analytical choices. Understanding which factors are most important to explaining the difference between immunopeptidomics and conventional shotgun proteomics may require the development of more sensitive proteomic techniques for identifying low-abundance and short-lived microproteins in the cell.

Why do several studies report low-quality spectra despite controlling FDR at 1%? Most of the studies we evaluated control only the proteome-wide FDR instead of controlling FDR for unannotated peptides or proteins specifically (Table 1).^17,23,59^ Since the proteome-wide FDR does not imply any particular FDR among unannotated proteins^17,23^, it does not imply high confidence in the unannotated list specifically. In a theoretical example experiment in which 1 million PSMs, 50,000 peptides and 10,000 proteins pass threshold, a 1% FDR corresponds to 10,000 incorrect PSMs, 500 incorrect peptides, or 100 incorrect proteins. If the analysis purports to detect 50 sORFs, the default assumption has to be that these are mostly part of the population of incorrect identifications until very carefully scrutinized. Studies that controlled FDR for unannotated proteins in a class-specific manner, such as Chong et al. 2020^42^ and Ouspenskaia et al. 2022^29^, scored high in our evaluations. We recommend that studies of the unannotated proteome report local or class-specific unannotated FDRs instead of, or in addition to, whole proteome FDRs, so that confidence in the list of reported unannotated proteins can itself be evaluated. To facilitate future work on the detection of unannotated microproteins by MS-based proteomics, we developed a set of guidelines based on our findings (brief advice in Box 1, detailed guidelines in Appendix 2). The guidelines in Appendix 2 are an extension of the Human Proteome Project Mass Spectrometry Data Interpretation Guidelines 3.0.^27^

### Box 1 Advice for detection of novel microproteins using mass spectrometry-based proteomics

- Ensure peptides appearing to support a novel protein detection uniquely support that protein:

- Conduct a search using tools such as ProteoMapper^45^ or PepQuery^60^ to exclude peptides with possible matches to canonical proteins, including post- and co-translational modifications and common genetic variants. When possible, construct a sample-specific protein database that accounts for genotype. Do not assume a canonical protein is absent from the sample solely on the basis of gene transcription or translation evidence.
- Consider whether the peptide may come from a previously unannotated isoform of a known protein-coding gene, as gene annotation databases do not comprehensively capture all transcript diversity. Ideally, integrate short- or long-read transcriptomics data to determine whether the evidence supports an unannotated alternative transcript or splicing event that could explain the observed translation.
- Pseudogene annotations can significantly impact microprotein discovery. Always check whether the peptide overlaps with a known pseudogene locus from either the Ensembl-GENCODE or RefSeq catalog.
- Ensure that the PSMs used to support a novel protein detection are high quality:

- Among PSMs that score highly in a search engine, spectra match quality can be further supported by comparison to experimental spectra generated from synthetized peptides, comparison to in silico fragmentation spectra generated by methods such as Prosit^61^ or MS2PIP,^48^ and machine learning rescoring using approaches such as Oktoberfest^50^ or MS2Rescore.^62^
- Manual evaluation of a representative subset of PSMs is important to ensure reported detections are supported by high quality evidence.
- To accurately convey confidence in the list of unannotated protein detections, report local FDRs or FDRs specific to the list of unannotated proteins instead of or in addition to proteome-wide global FDR. The less stringent the FDR threshold used, the more it is necessary to examine candidates further to ensure they are correct.
- Make the MS data available in a public data repository. Report universal spectrum identifiers (USIs)^63^ for all spectra supporting discovery of a novel protein.

It is important to note that false positives can occur across the full range of PSM quality; a low-quality spectrum does not prove that a claimed detection is a false positive; nor is a high-quality spectrum conclusive evidence of detection. The gold standard for rigorous MS-based proteomics data validation requires demonstration that a synthetic peptide generates the observed spectrum and is retained on the liquid chromatography column to the same extent as the originally detected peptide, and that the endogenous spectrum is eliminated when the ORF is disabled genetically. Supporting evidence for the biological significance of a protein with inconclusive MS support can also come from outside proteomics, such as by demonstrating the evolutionary conservation of its amino acid sequence or reporting phenotypic impacts upon genetic perturbations.^23,53^

The thousands of sORFs identified by Ribo-Seq experiments suggest a massive potential for undiscovered microproteins of biomedical relevance, even at low proteomic validation rates. While our community assessment found relatively low proteomic support for these microproteins in the datasets generated by the pioneering studies we analyzed, this finding should not be interpreted to mean that only few sORF-encoded proteins are present in the cell. There are major technical limitations in the ability of proteomic experiments to find short and low-abundance proteins^16,23,25^, and the microproteins field is still in its infancy. The extent to which sORFs encode stable functional proteins thus remains an open question. To answer it, we will need to expand the limits of protein detectability through further methodological developments, including but not limited to improving the sensitivity of MS instruments. We hope the dataset of 406 manually curated PSMs generated here will prove useful for benchmarking much-needed new data analysis tools and pipelines for unannotated microprotein detection by MS (Supplementary Table 4).

## Methods

### Study selection

We conducted a search for all studies published in the 2019-2022 period that attempted to detect unannotated proteins using shotgun proteomics. For each study, we obtained information on the PSMs claimed to support each reported unannotated detection (Supplementary Table 1). For each PSM, we collected the information needed to construct a universal spectrum identifier (USI)^63^ so the PSM could be visualized. Where possible, we obtained the PSM data from the supplementary information provided with the study; otherwise, we attempted to obtain them from the study authors. The sources of data for each study are given in Supplementary Table 2. The authors of one study (Cai et al. 2021)^64^ were unable to provide the necessary data so this study was not evaluated.

The set of “unannotated” proteins depends on the annotation database used; the proteins included in our analysis followed the definition used in each study. Unannotated proteoforms of annotated proteins were not included.

### ProteoMapper analysis

All reported unannotated peptides were submitted to the ProteoMapper online tool^45^ in July 2023 using default settings. For each peptide, ProteoMapper returns a list of matches to known or predicted proteins, accounting for neXtProt^46^ amino acid variants. We determined whether each peptide mapped to a human annotated protein according to the 2023 build of the PeptideAtlas database^65^ and whether each peptide mapped to a protein present in the 2016 version of UniProtKB/Swiss-Prot.^18^ Any peptide that mapped to a core canonical PeptideAtlas protein on ProteoMapper was not passed on for manual evaluation, even if it differed from the reference sequence by multiple neXtProt amino acid variants.

### Manual evaluation of PSM quality

PSMs for each study were evaluated by a group of six expert evaluators. Each evaluator rated a random sample of PSMs from each study. A total of 424 PSMs from 12 studies were given for evaluation, out of which 406 were given ratings, as a few PSMs could not be displayed from the input USI. Out of the 406 PSMs evaluated, 155 were evaluated by two evaluators each to enable determination of the overall consistency between evaluators. Evaluations were done by visual inspection of the PSM using the ProteomeCentral USI web application (https://proteomecentral.proteomexchange.org/usi/) in May to June 2023. The evaluators were told to use no other information except the PSM as displayed on the USI application. A common set of 10 negative control PSMs was given to each evaluator; the evaluators were not informed of the existence of these controls. These negative controls consisted of high-scoring decoy-spectrum matches manually selected from among the strongest 30 decoy-spectrum matches in Duffy et al. 2022.^34^ Each PSM was rated on a scale of 1-5; the rating scale is given in Appendix 1.

### Comparing manual evaluations to spectral prediction machine learning methods

Spectra were predicted for each manually evaluated peptide sequence annotated to the set of experimental spectra using the open-source spectral library prediction pipeline Oktoberfest.^50^ Multiple predicted spectra were generated for each peptide at various collision energies (CE = 25, 30, 35 and 40) and using 4 different intensity models (Prosit 2020 intensity HCD^61^, Prosit 2020 intensity CID, Prosit 2020 intensity TMT, AlphaPept ms2 generic)^61,66–68^. Only methionine oxidation, cysteine carbamidomethylation, and TMT6plex modifications were considered in the spectral predictions; peptides with other modifications were excluded for this analysis. MSP spectral library files output by Oktoberfest were then converted to MGF formatted spectra. Internal python scripts compared the experimental spectra vs. the predicted spectra by calculating spectral angles (SA) between each spectral pair. Similarity was ranked as being high if SA ≤ 20^⁰^, moderate if SA between 20^⁰^ - 45^⁰^, poor if SA between 45^⁰^ - 70^⁰^, and terrible if SA > 70^⁰^. The script further generated mirrored plots for each spectral pair and annotated peptide fragment ions. These spectral angles were then compared to the manual ratings for each PSM given by the evaluators.

### Predicting HLA binding for immunopeptides supporting unannotated protein detections

For each evaluated immunopeptide from Ouspenskaia et al. 2021, Martinez et al. 2020, or Chong et al. 2020 used to support an unannotated protein detection, the HLA alleles for the cell type used in the experiment producing the peptide was found in the supplemental data of the study. NetMHC 4.0 was then used to predict binding of the peptide to the HLA-A, HLA-B, and HLA-C allele if the allele was available in NetMHC 4.0. A peptide was classified as being HLA-binding if it met the default criteria for being a weak (% rank < 2%) or strong (% rank < 0.5%) binder in NetMHC 4.0.

### Relating ORF properties to the probability of detection

The coordinates of each ORF with an evaluated peptide were taken from the supplementary data of each study and the ORF length determined. All ORF coordinates were converted to hg38 coordinates using LiftOver. ORFs from Chen et al. 2020^41^, Chong et al. 2020^42^, Cao et al. 2022^30^, and Lu et al. 2019^44^ were not considered because we were not able to identify the ORF coordinates from supplementary data files. To assess translation levels, we aggregated Ribo-Seq data from 109 studies (Supplementary Tables 5-6) using the following procedure. Transcriptomes from MiTranscriptome^69^, FANTOM5 robust set^69^, CHESS^70^, RNA Atlas^71^, and Ensembl version 108 were merged using Stringtie^72^ version 2.2.1with Ensembl version 108 as the reference annotation (-G parameter). MiTranscriptome and FANTOM5 coordinates were lifted over from hg19 to hg38 prior to merging. Adapters in each ribo-seq run were removed with TrimGalore version 0.6.7 using default options. Trimmed Ribo-seq reads were then mapped to the merged transcriptome using STAR^73^ version STAR-2.7.10b using the parameters-- outSAMtype BAM Unsorted --outFilterMismatchNmax 2 --outFilterMultimapNmax 1 --outSAMattributes Standard. The iRibo program^54^ was then used to aggregate the mapped reads from all studies and assign counts of ribosome P-sites to each position of each analyzed ORF.

## Data Availability Statement

All data analyzed are available at: https://github.com/CarvunisLab/MSCommunityAssessment

## Code Availability Statement

All code used for analyses are available at: https://github.com/CarvunisLab/MSCommunityAssessment

## Supplementary Tables

**Supplementary Table 1: Properties of all PSMs reported to support unannotated protein detections that were considered in this study.**

**Supplementary Table 2: Source of PSM data for each analyzed study**

**Supplementary Table 3: Studies reporting detection of each peptide.**

**Supplementary Table 4: Evaluator ratings for each evaluated PSM.**

**Supplementary Table 5: NetMHC binding prediction for evaluated immunopeptides**

**Supplementary Table 6: Ribo-seq studies analyzed**

**Supplementary Table 7: Translation levels for analyzed ORFs**

## Author contributions

Conceptualization: Aaron Wacholder, Eric W. Deutsch, Sebastiaan van Heesch, John R. Prensner, Thomas F. Martinez, Marie A. Brunet, Jana Schulz, Jorge Ruiz-Orera, Jonathan M. Mudge, Sarah A. Slavoff, Anne-Ruxandra Carvunis

Methodology: Aaron Wacholder, Anne-Ruxandra Carvunis, Eric Deutsch

Formal analysis: Aaron Wacholder, Jiwon Lee, Sebastien Leblanc, James C. Wright, Leron W. Kok, Jip T. van Dinter

Investigation: Aaron Wacholder, Eric W. Deutsch, John R. Prensner, Thomas F. Martinez, Marie A. Brunet, Jorge Ruiz-Orera, Jonathan M. Mudge, Sarah A. Slavoff

Resources: Eric W. Deutsch

Data Curation: Ihor Arefiev, Francis Bourassa, Kevin Cao, Ayodya H Jayatissa, Kevin Jiang, Felix-Antoine Trifiro, Eric Deutsch

Writing - Original Draft: Aaron Wacholder

Writing - Review & Editing: Sebastiaan van Heesch, Leron W. Kok, Jip T. van Dinter, Ivo Fierro-Monti, Eric W. Deutsch, Michal Bassani-Sternberg, Sonia Chothani, Juan Antonio Vizcaíno, Jyoti S. Choudhary, Marie A. Brunet, Xavier Roucou, Jonathan M. Mudge, John R. Prensner, Pavel V. Baranov, Jorge Ruiz-Orera, Norbert Hubner, Sarah A. Slavoff, Thomas F. Martinez, Annelies Bogaert, Daria Fijalkowska, Kris Gevaert, Robert L. Moritz, Anne-Ruxandra Carvunis Visualization: Aaron Wacholder

Project administration: Aaron Wacholder and Anne-Ruxandra Carvunis Supervision: Anne-Ruxandra Carvunis

## Acknowledgments

This work was supported in part by a Research Grant from HSFP awarded to AR-C: https://doi.org/10.52044/HFSP.RGP0042023.pc.gr.168590. M.B.-S. is supported by the Ludwig Institute for Cancer Research, by grants KFS-4680-02-2019 and KFS-5637-08-2022 from the Swiss Cancer Research Foundation (M.B.-S.), the Swiss National Science Foundation PRIMA grant PR00P3_193079 (M.B.-S.) and the Swiss Bridge Foundation Award (M.B.S). J.A.V. is supported by funding from Wellcome [grant number 223745/Z/21/Z], and from EMBL core funding. J.S.C. acknowledges funding from the Wellcome Trust [223745/Z/21/Z] and from the ICR core funding. M.A.B is supported by a Junior 1 career award from the Fonds de Recherche du Quebec - Sante (FRQS). F.B. is supported by a FRQS scholarship. F.A.T. is supported by a FRQS scholarship. I.A. is supported by a FRQS scholarship. X. R. is supported by the Canadian Institutes for Health Research (CIHR) (Grant No. PJT-175322), and Canada Research Chair in Functional Proteomics and Discovery of Novel Proteins. J. M. M. is supported by the Wellcome Trust (108749/Z/15/Z), the National Human Genome Research Institute (NHGRI) of the U.S. National Institutes of Health (NIH) under award number (2U41HG007234), and the European Molecular Biology Laboratory (EMBL). The content is solely the responsibility of the authors and does not necessarily represent the official views of the National Institutes of Health. Ensembl is a registered trademark of EMBL. K. J. is supported in part by a NIH Chemical Biology training grant (T32 GM149444). J.R.P. acknowledges funding from the National Institutes of Health / National Cancer Institute [K08-CA263552-01A1]; the V Foundation for Cancer Research [V2024-013]; Hyundai Hope on Wheels Foundation; the Yuvaan Tiwari Foundation; DIPG/DMG Research Funding Alliance; Tough2gether Foundation; CureSearch Foundation; Morgan Adams Foundation; ChadTough Defeat DIPG Foundation; Book for Hope Foundation; Curing Kids Cancer Foundation [20-3388093], and the Andrew McDonough B+ Foundation [1185689]. J.R.P. is the Ben and Catherine Ivy Foundation Clinical Investigator of the Damon Runyon Cancer Research Foundation [CI-127-24]. S. A. S. is supported by the Paul G. Allen Frontiers Group Distinguished Investigator Award. This work was funded in part by the National Institutes of Health grants R24 GM148372 (E.W.D.), R01 GM087221 (E.W.D., R.L.M.), S10 OD026936 (R.L.M.), and by National Science Foundation grants DBI-2324882 (E.W.D.) DBI-1933311 (E.W.D.), and MRI-1920268 (R.L.M.). N.H. was supported by a grant from the Leducq Foundation, an ERC Advanced Grant under the European Union Horizon 2020 Research and Innovation Program (AdG788970), a British Heart Foundation and a Deutsches Zentrum für Herz-Kreislauf-Forschung grant (BHF/DZHK: SP/19/1/34461), by German Research Foundation - DFG (CRC/SFB-1470 – B03), and in part by a grant from the Chan Zuckerberg Foundation (2019-202666). J.C.W. acknowledges the support of The Institute of Cancer Research and funding from Wellcome [grant numbers 208391/Z/17/Z, 223745/Z/21/Z]. S.L. is supported by Canadian Institutes for Health Research (CIHR) (Grant No. PJT-175322), and Canada Research Chair in Functional Proteomics and Discovery of Novel Proteins. P.V.B. is supported by Taighde Éireann – Research Ireland under Grant number [20/FFP-A/8929]. K.G. was supported by The Research Foundation—Flanders (FWO), project number G008018N. S.v.H. acknowledges funding from Fonds Cancers (FOCA, Belgium), Stichting Reggeborgh (the Netherlands), and Villa Joep. This publication is part of the project “Evolutionarily young microproteins in childhood brain cancer” (with project number VI.Vidi.223.022 of the research programme NWO talent programme Vidi, which is (partly) financed by the Dutch Research Council (NWO), awarded to S.v.H. Research reported in this publication was supported by Oncode Accelerator, a Dutch National Growth Fund project under grant number NGFOP2201, awarded to S.v.H. I.F-M. financial support was received from the European Union’s Horizon 2020 research and innovation programme under the Marie Sklodowska-Curie grant agreement No. 945405 (ARISE programme). S.C. is funded by Singapore Ministry of Health’s National Medical Research Council under OF-YIRG (OFYIRG23jan-0034). We are grateful for helpful feedback from Aviv Regev, Travis Law, Tamara Ouspenskaia, Karl Clauser, Susan Klaeger, Catherine J. Wu, Owen Rackham, Gong Zhang, Michelle Magrane, Erin Duffy, Brian Kalish, and Michael E. Greenberg.

## Appendix 1 PSM rating scheme and examples

The following scheme was given to evaluators as a basis for rating each PSM. Below, example PSMs are given together with an explanation for their rating provided by an evaluator. Each PSM can be visualized on ProteomeCentral (https://proteomecentral.proteomexchange.org/usi/) using the USI.

5 - Excellent. A very good match that shows peaks for nearly every residue except perhaps b1 (and thus the order of the two utmost N-terminal ions is unclear) and is not contaminated by a cofragmented precursor. Really solid evidence for the peptide. Example: mzspec:PXD021482:20200724_cell_10:scan:6083:APQSPGPAPPPASSGR/2

4 - Good enough even for an extraordinary detection. Perhaps one or two peaks missing, perhaps some contamination, but seems like a good match. Example: mzspec:PXD014058:20181120_HCT116_P-ACN_up_14:scan:35747:GGQSLPTTMWSPVK/2

3 - Not good enough. It might be right. But it might well be something else fairly close. Incomplete coverage of the residues. This PSM would be fine if it were a common albumin peptide, but the bar for a non-canonical ORF is higher. Examples:

mzspec:PXD020079:20180504_QEh1_LC1_QC_JMI_HLAIp_HROG17_2_R2:scan:6918:AVAGSRGDKSLR/3

mzspec:PXD010154:01698_A02_P018021_S00_N09_R1:scan:11204:ASEIQSTGGQRDPQPER/3

mzspec:PXD004894:20141216_QEp7_MiBa_SA_HLA-I-p_MMf_6_2:scan:29039:KPRLPIYGL/2

mzspec:MSV000080527:M20151203_HLA_A2402_75millionceq_biorep1_techrep1:scan:46144:YSLSLQIL F/2

2 - Wrong with a high quality spectrum. Clearly not the correct interpretation, although perhaps close, but this spectrum probably has a good alternative explanation. Usually, some high unexplained peaks near but not exactly at where there ought to be a peak if the interpretation were correct. Example:

mzspec:PXD004894:20141215_QEp7_MiBa_SA_HLA-I-p_MMf_15_1:scan:34668:MPRMALVYHTA/3

1 - Poor quality spectrum. Not enough information to be sure about any id. Or perhaps a clearly blended spectrum where it’s hard to be sure of any id. Example:

mzspec:PXD019643:170421_AM_AUT01-DN14_BoneMarrow_W6-32_10__DDA_2_400- 650mz_msms23_standard:scan:26200:SVWLSPPPA/2

**Example PSMs with consistent results from the evaluators**

Example of a 5 star rating: Both reviewers gave it a 5.

mzspec:PXD999953:09CPTAC_UCEC_W_PNNL_20180222_B3S1_f03:scan:33089:[TMT6plex]-NDDIPEQDSLGLSNLQK[TMT6plex]/2

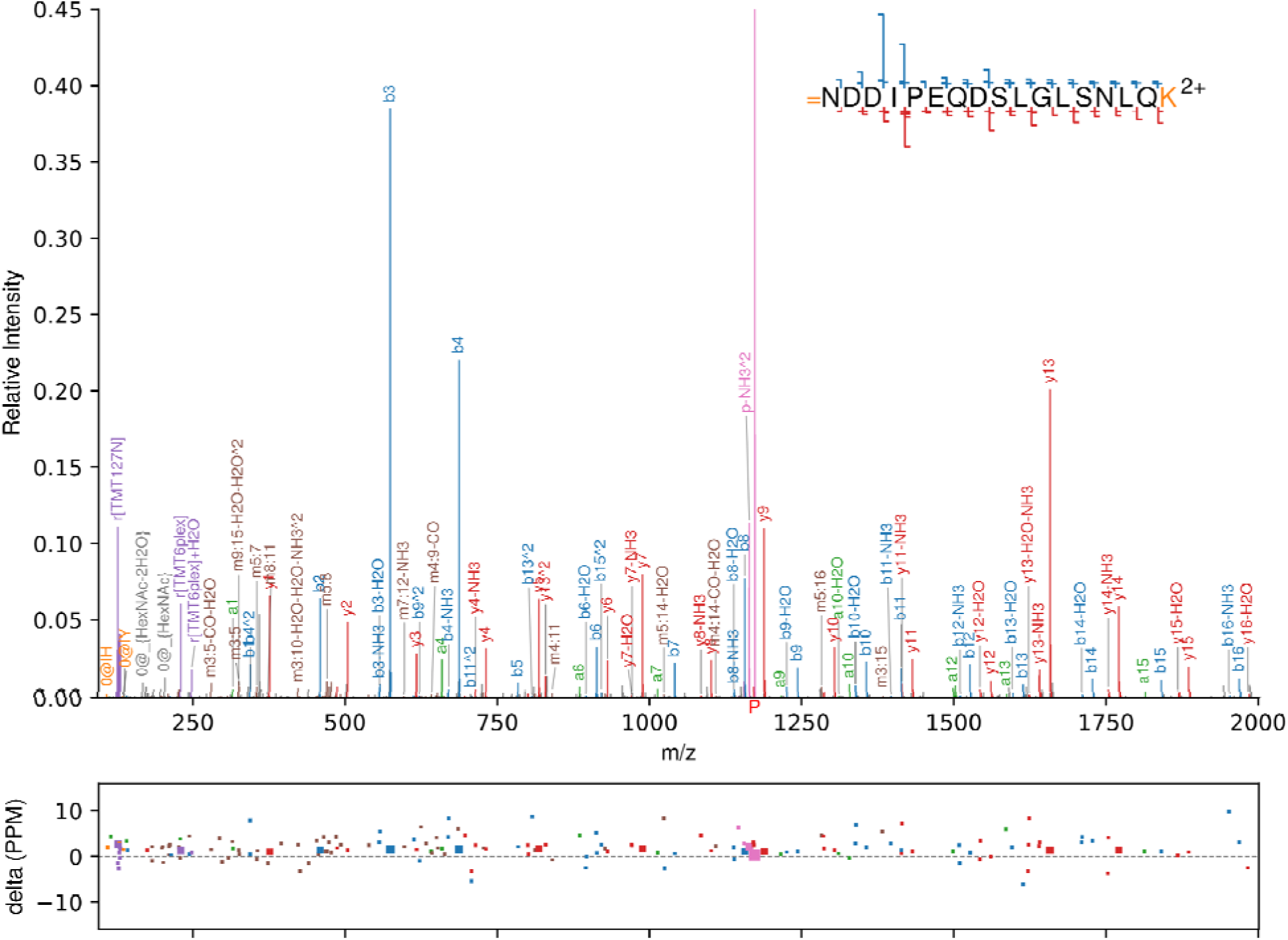

This PSM presents a very strong match with full coverage from both ends. This 64 amino acid protein was highlighted in Kim et al. 2014. It is currently in UniProtKB/TrEMBL as Q9HB66 but not in UniProtKB/Swiss-Prot.

Example of a 4 star rating: Both reviewers gave it a 4.

mzspec:PXD026880:VOT16-2132:scan:3865:AAPELGPGATIEAGAAR/2

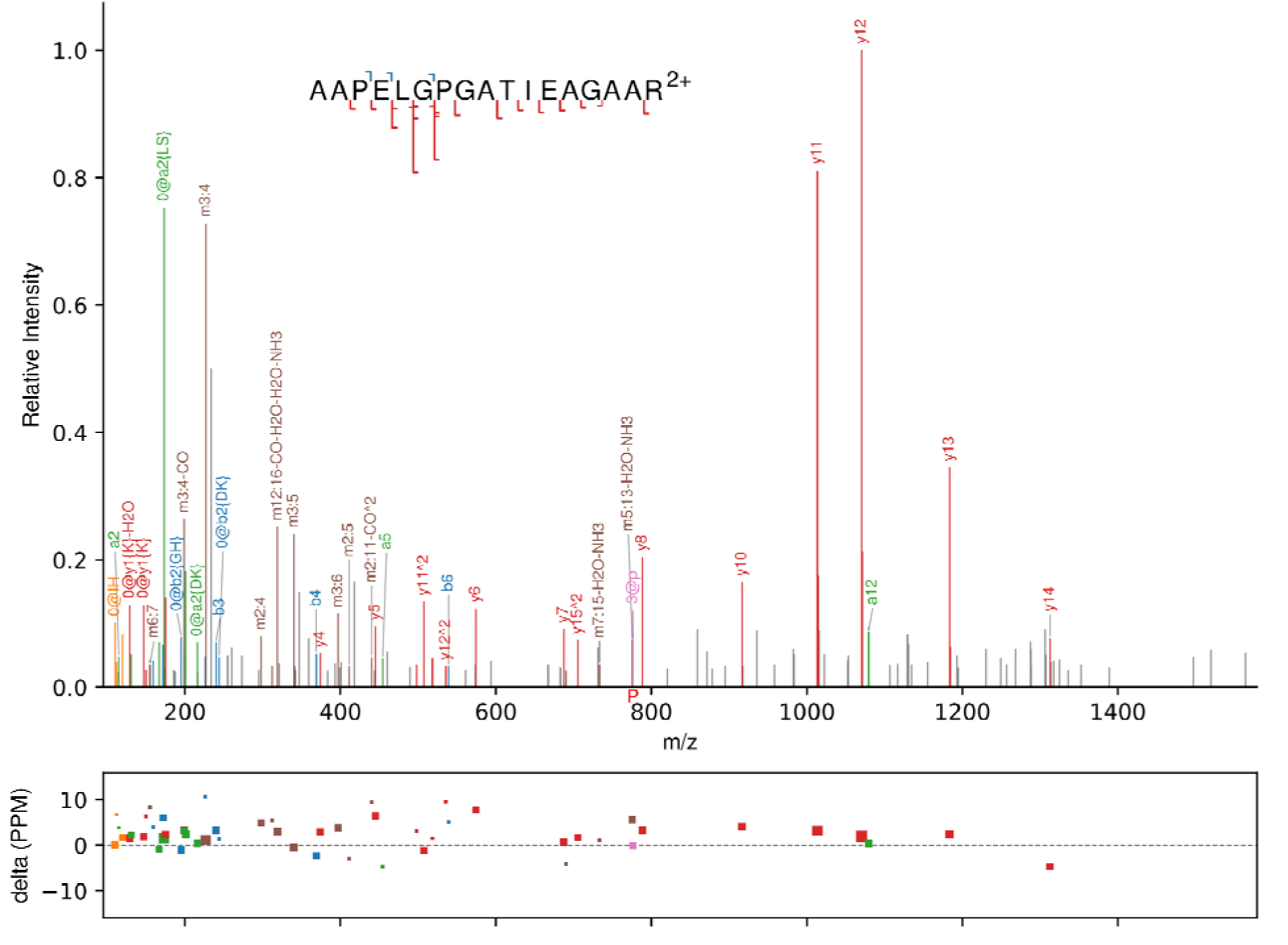

This PSM provides nearly complete coverage in y ions, although there are some gaps. The b ions are very weak, but that is not surprising given the sequence. Signal to noise (as estimated by the ratio of the tallest to smallest peak) is decent, but weaker than the PSMs rated 5. The precursor m/z value is exactly as expected.

There are a few major peaks that are not easily explained except by a y-ion series of a contaminating peptide ending in LSK, explaining peaks at 147.1311, 235.1455, and 347.2301. This reduces confidence slightly. For these reasons, this PSM does not rate a 5, but is good evidence for the peptide.

Example of a 3 star rating: All 3 reviewers gave it a 3.

mzspec:PXD026880:VOT16-2132:scan:9332:EPAGPSLNPTLAATAALIPLHR/3

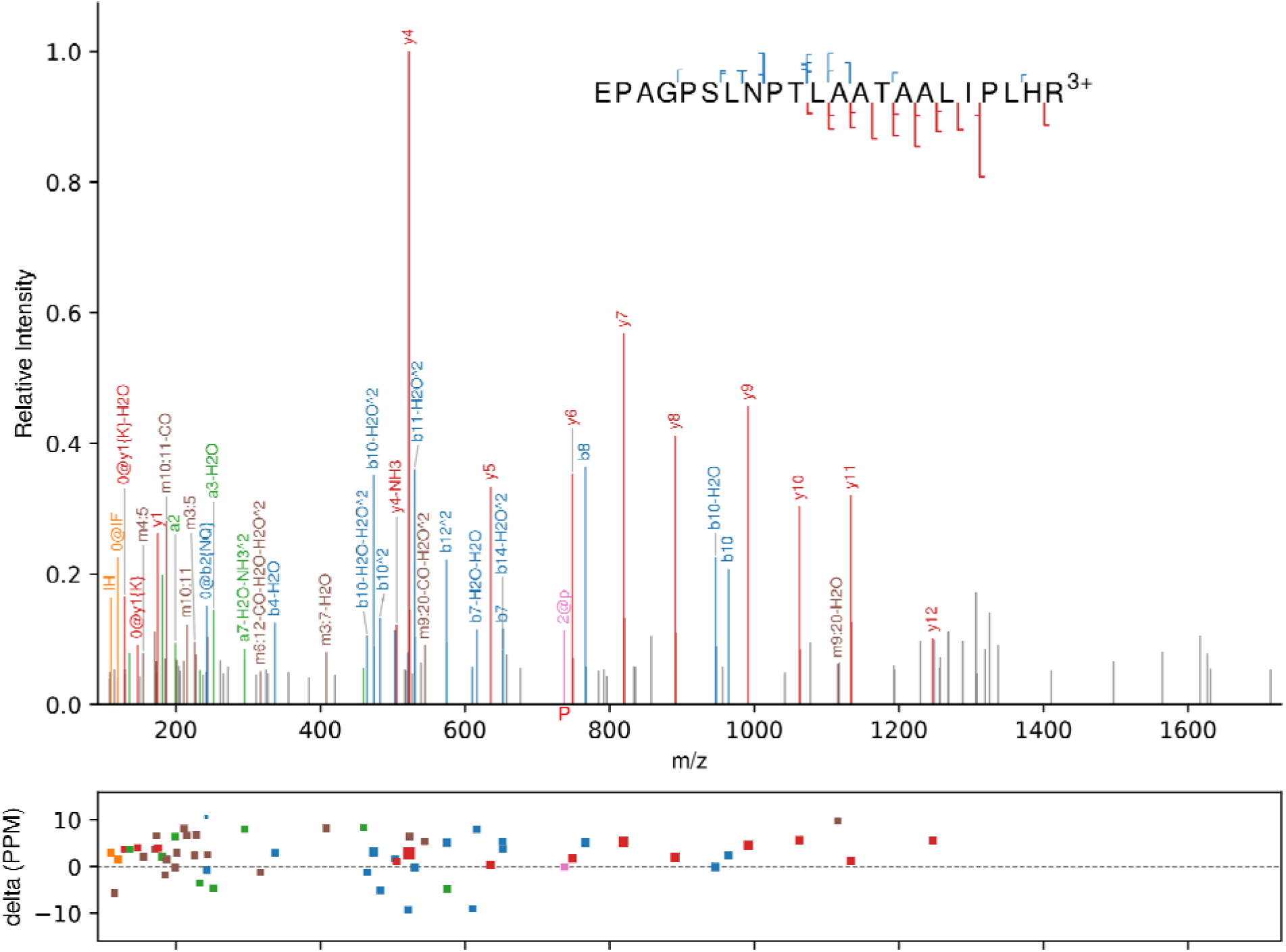

There is good evidence for the second half of the peptide, but evidence is almost entirely lacking for the first half of the peptide. The peptide sequence is likely at least partially correct but may have a different N terminus. If this were an annotated protein, it would be sufficient, but not for a claim of a novel protein.

Example of a 2 star rating: Both reviewers gave it a 2.

mzspec:PXD014031:20190104_QX4_AnBr_SA_IPSC_Peptidome_Fraction_12:scan:94981:SLEGLIPSSSVVGK/2

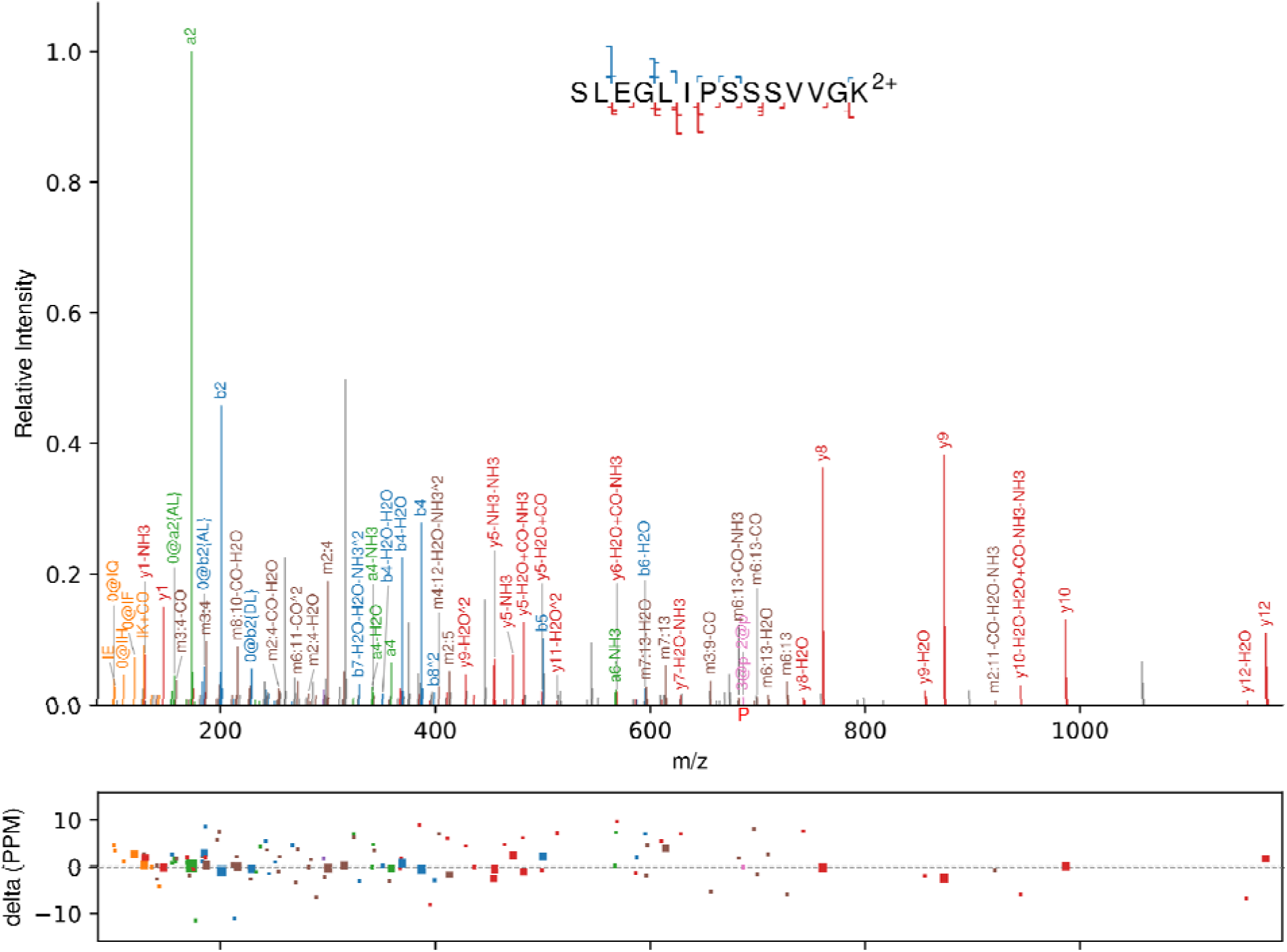

This is a high-signal-to-noise spectrum, and it is clear why the search engines gave it a high score, but there are several major unexplained peaks in regions where we expect good peaks. Due to the lysine on the C terminus and no other K, R, or H residues in the sequence, the lack of any y2-y7 fragment identifications coupled many unexplained peaks below 600 m/z is a strong indicator of an incorrect identification.

This spectrum is most likely derived from a different peptide than claimed. Manual interpretation of the spectrum reveals a far better matched peptide: mzspec:PXD014031:20190104_QX4_AnBr_SA_IPSC_Peptidome_Fraction_12:scan:94981:SLDALISQVADLK/2

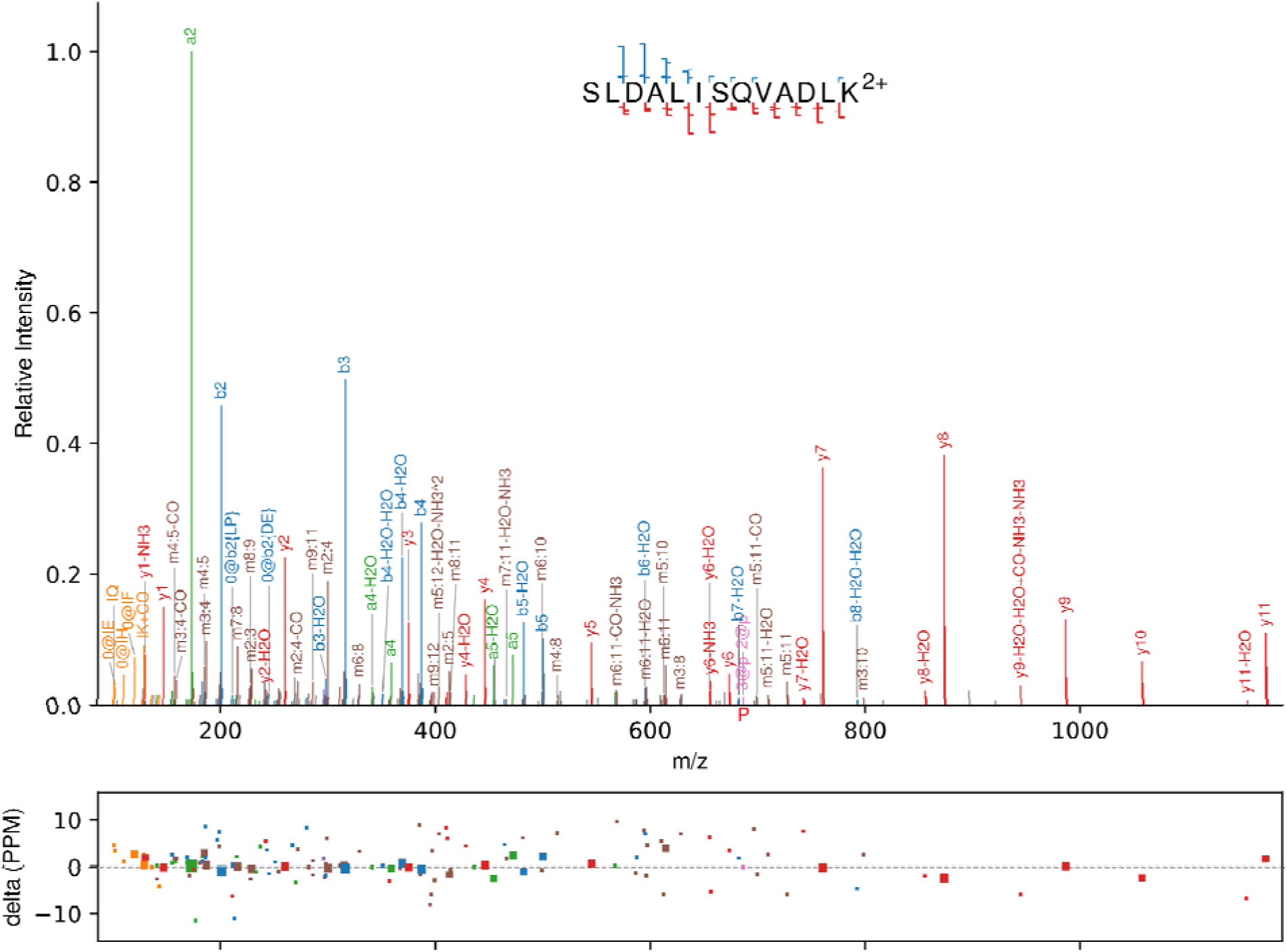

This is a much better PSM for the spectrum, matching the very well detected protein Q96G25 (Mediator of RNA polymerase II transcription subunit 8). This semi-tryptic peptide has 54 PSMs in the 2024 human PeptideAtlas. However, this peptide is not in the search space of a fully tryptic search. Only a semi-tryptic search will find it. A fully tryptic search may lead to false positives if something else in the search space is similar.

Example of a 1 star rating: Both reviewers gave this spectrum a rating of 1.

mzspec:PXD006675:20160721_QEp2_SoDo_SA_LC12-13_RV8-frac2:scan:65697:LLQTLAKNFIGK/3

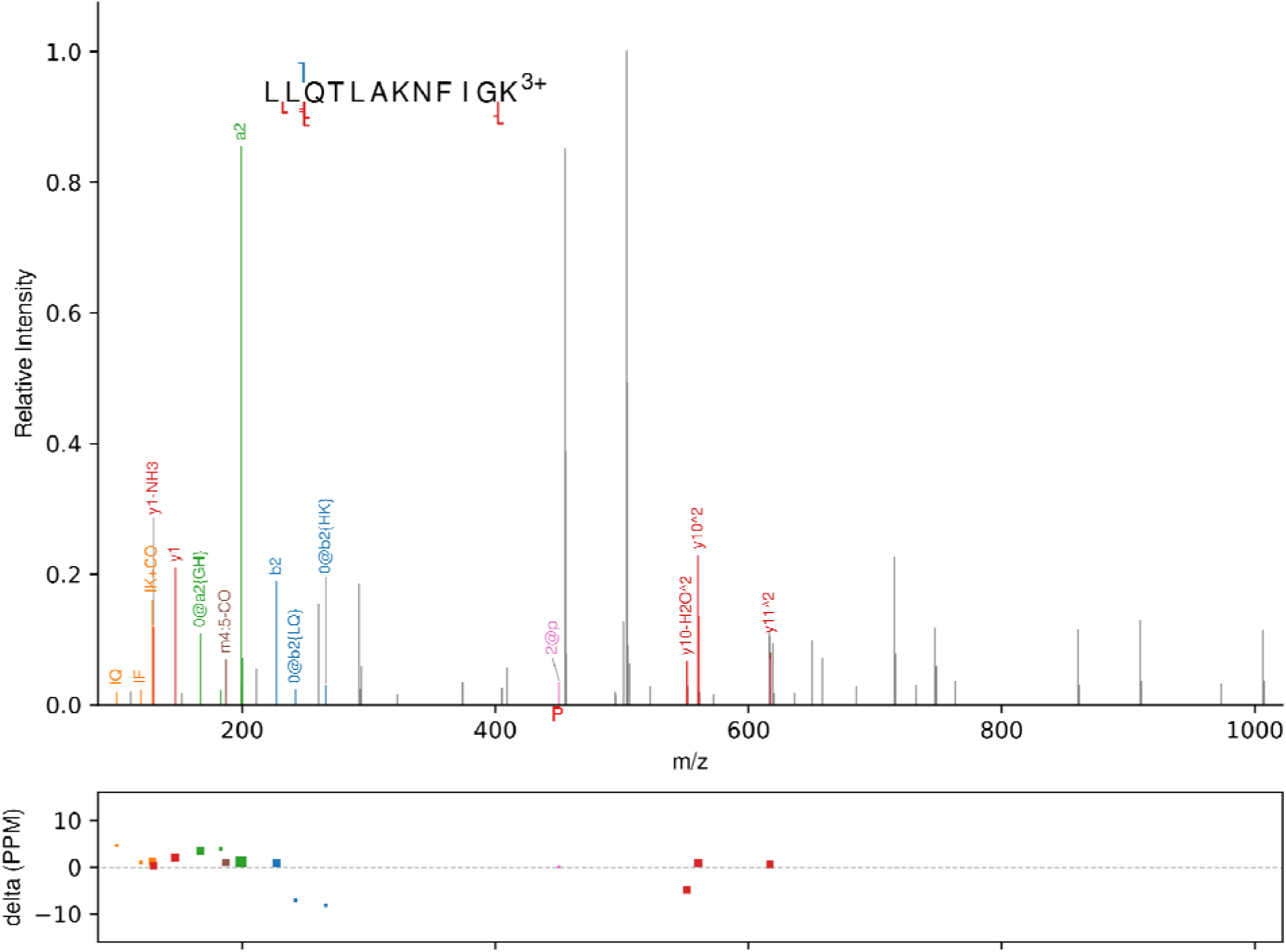

This is a low signal-to-noise spectrum that is very poor evidence for the claimed peptide and may be too low quality to confidently identify any peptide.

Manual interpretation of the spectrum reveals a peptide that fits far better:

mzspec:PXD006675:20160721_QEp2_SoDo_SA_LC12-13_RV8-frac2:scan:65697:LLLPHGVDQLLK/3 explaining most peaks in the spectrum, but it is unclear where that peptide might derive from, as it does not map to known proteins, and there are still gaps in coverage.

## Appendix 2 Mass Spectrometry Microprotein Detection Guidelines

Mass Spectrometry Microprotein Detection Guidelines Version 0.2.0 – June 17, 2025

| General guidelines for all manuscripts: |  |  |
| --- | --- | --- |
| √ | Loc | 1. Complete this MS Microprotein Detection Guidelines checklist |
| Data deposition and reference proteome guidelines |  |  |
|  |  | 2a. Deposit all MS proteomics data to a ProteomeXchange repository as a “complete” submission. |
|  |  | 2b. Include analysis reference files (search database, spectral library, transition list, etc.) in submission. |
|  |  | 2c. Provide the PXD identifier(s) in the manuscript abstract. |
|  |  | 2d. Provide the reviewer login credentials if the dataset is not yet public. |
|  |  | 3. Use the most recent version of the HPP Target List plus UniProtKB reviewed isoforms plus contaminants as the reference canonical proteome for all informatics analyses. |
| FDR-related guidelines |  |  |
|  |  | 4a. Describe in detail the calculation of FDRs at the PSM, peptide, and protein levels. |
|  |  | 4b. Report the PSM-, peptide-, and protein-level FDR values along with the total number of expected false positives at each level, using precision appropriate to the uncertainty in computed FDR. |
|  |  | 4c. Present large-scale results thresholded at equal to or lower than 1% protein level global FDR, but not more than 10% local FDR. |
|  |  | 4d. If any large-scale datasets are individually thresholded and then combined, calculate the new, higher peptide- and protein-level FDRs for the combined result. |
| Guidelines for claims of new protein detections |  |  |
|  |  | 5a. If using DDA mass spectrometry for such claims, present high mass-accuracy, high-SNR, and clearly annotated spectra to ensure PSMs are correct. Scrutinize spectra for missing and extra peaks. |
|  |  | 5b. Present high mass-accuracy, high-SNR, clearly annotated spectra of synthetic peptides that match the spectra supporting the claims. Compute normalized backbone ion dot products between target spectra and synthetic peptide spectra, as well as between target spectra and predicted spectra. |
|  |  | 5c. Provide Universal Spectrum Identifiers (USIs) for all natural, synthetic, and predicted peptide-spectrum matches that support such claims, ideally as a supplementary data table. |
|  |  | 6. If using SRM verification for such claims, present target traces alongside synthetic heavy-labeled peptide traces, demonstrating co-elution and closely matching fragment mass intensity patterns. |
|  |  | 7. If using DIA MS, then, if the data are analyzed with XICs, apply the above SRM guidelines (6); if the data are analyzed by extracting deconvoluted spectra, apply the above DDA guidelines 5a-5c. |
|  |  | 8. Even when very high confidence peptide identifications are demonstrated, consider alternate mappings of each peptide to canonical proteins. Consider isobaric sequence/mass modification variants, all known SAAVs, and unreported SAAVs. |
|  |  | 9. Support Tier 1 (1A digest, 1B HLA) claims via two or more distinct uniquely-mapping, non- |
| | | nested peptide sequences of length $\geq 9$ amino acids with the above evidence. When 2 peptides overlap, the total extent must be $\geq 18$ amino acids. 100% coverage, except for an initiating methionine, may be considered Tier 1 evidence. Tier 2A (digest) or Tier 2B (HLA) evidence may constitute just a single uniquely-mapping peptide of length $\geq 9$ amino acids. Tier 3 evidence (not meeting Tier 2A or 2B evidence) may be discussed but is generally not considered sufficient for a claim of detection. |

See extended description for each of the above items on pages 2 - 4 below.

**Extended Detail on Checklist items:**

The following pages provide some additional detail on the intentionally terse one-page checklist. Users should read these extended descriptions before using the checklist.

1. **Complete this MS Data Protein Detection Guidelines checklist.**
2. **Guidelines for data repository deposition**

a. **Deposit all MS proteomics data to a ProteomeXchange repository as a complete submission.** All depositions should be “Complete” submissions instead of “Partial” submissions. ProteomeXchange deposition should be completed prior to submission of the manuscript to the journal. Synthetic peptide MS runs should also be deposited and clearly marked as such.
b. **Include analysis reference files (search database, spectral library, transition list, etc.) in submission.** Include all supplemental data files used in the analysis. Include software parameter files if relevant.
c. **Provide the PXD identifier(s) in the manuscript abstract.**
d. **Provide the reviewer login credentials if the dataset is not yet public.** Reviewer login information at the repository should be provided in the manuscript if the dataset is not already publicly released.
3. **Use the most recent version of the HPP Target List plus UniProtKB reviewed isoforms plus contaminants as the reference canonical proteome for all informatics analyses.** Informatics analysis should always be presented in comparison with the most recent canonical proteome references, rather than older versions thereof. The most recent versions of the HPP Target List, UniProtKB isoforms, and contaminants may be found at the HPP Portal and PeptideAtlas.
4. **FDR-related guidelines**

a. **Describe in detail the calculation of FDRs at the PSM, peptide, and protein levels.** Describe which tools are used to estimate the false discovery rate (FDR) at the peptide-spectrum-match (PSM) level, at the distinct peptide sequence level, and at the protein level. Briefly describe the approach and what assumptions are made or implied, and any corrections for the fraction of the proteome covered. If you use novel or uncommon tools and criteria, compare your results with results with tools that are widely used in the community.
b. **Report the PSM-, peptide-, and protein-level FDR values along with the total number of expected false positives at each level, using precision appropriate to the uncertainty in computed FDR.** Report the actual numbers of true positives and false positives at each level based on the thresholds used. Do not report the FDR with many significant digits since all current FDR calculation methods have substantial uncertainties.
c. **Present large-scale results thresholded at equal to or lower than 1% protein-level global FDR, but not more than 10% local FDR. The 1% is somewhat arbitrary but well accepted and remains set as the upper limit.** As is frequently the case with datasets from modern instrumentation, a local FDR of 10% is reached before a 1% global FDR is reached, and then 10% local FDR threshold or less should be used instead. The common mistake of thresholding at a specific FDR and then proceeding as if all surviving results are correct, no matter how surprising, must be avoided.
d. **If any large-scale datasets are individually thresholded and then combined, calculate the new, higher peptide- and protein-level FDRs for the combined result.** When datasets are combined, the true positives will mostly overlap, while the false positives will be scattered randomly across the proteome and thus overlap far less. This means that the FDR will be higher in the combined dataset. Whereas the above guidelines apply generically to overall dataset analysis, the following guidelines apply specifically to the presentation of evidence of proteins that are not currently listed in the human reference proteome.
5. **Guidelines for data-dependent acquisition (DDA) MS datasets**

a. **If using DDA mass spectrometry for such claims, present high mass-accuracy, high signal-to-noise ratio (SNR), and clearly annotated spectra to ensure PSMs are correct. Scrutinize spectra for missing and extra peaks.** Annotated spectra (i.e., spectra with the matched peaks clearly labeled) must be provided in the supplementary material for the manuscript. While low mass-accuracy and low SNR spectra can still be useful for many experiments, they are not acceptable for claims of new protein detections. Time-of-flight, FT-ICR, Astral, and Orbitrap-type instruments are considered in these guidelines as having high mass accuracy (when properly calibrated) in these guidelines. The spectra should be examined closely to determine if there are peaks missing that should be expected, if there are peaks present that are unexplained, and if a small alteration of the putative sequence would yield a much better match. This may indicate a false positive of a kind that is not modeled well by decoys.
b. **Present high mass accuracy, high-SNR, clearly annotated spectra of synthetic peptides that match the spectra supporting the claims. Compute normalized backbone ion dot products between target spectra and synthetic spectra, as well as between target spectra and predicted spectra**. Synthetic peptides are powerful tools for determining the correct identification of spectra. For each PSM corresponding to claim of a new PE1 protein, compare that PSM with a synthetic peptide (or recombinant protein product) spectrum of the same ion. All the major ions should have closely matching intensities in both spectra. If generating new reference spectra, it is encouraged to use the same high mass-accuracy instrument to verify matching intensity patterns and retention times. Closely matching spectra of the same peptidoform ion (same modifications and charge) from SRMAtlas, ProteomeTools, or similar resources is acceptable. Predicted spectra may be used in addition to or even in place of synthetic reference spectra. Compute normalized (0 to 1) dot product based on all backbone ions (typically b & y) (omitting b1 and y1) within acquired m/z range between the target spectra and reference spectra and report the results.
c. **Provide Universal Spectrum Identifiers (USIs) for all natural and synthetic peptide spectra that support such claims, ideally as a supplementary data table.** The USI provides a mechanism to uniquely identify a spectrum being held up as evidence for an important claim. The USI will allow readers to access these important spectra in public data repositories in order to discuss correctness of the claims. See http://psidev.info/USI for more information.
6. **If using SRM verification for such claims, present target traces alongside synthetic heavy-labeled peptide traces, demonstrating co-elution and closely matching fragment mass intensity patterns.** All SRM runs performed must have spiked-in heavy labeled peptides corresponding to the putative identifications. The heavy-labeled peptides should be spiked in at an abundance similar to the target peptides so that minor impurities in the reference do not contribute to the target signal. Annotated chromatograms must be provided in the supplementary material of the manuscript. Solid peptide sequence evidence does not alter the uncertainties in matching that peptide uniquely to a protein (guideline 8). This guideline may also apply to PRM traces, although since PRM generates full MS/MS spectra, Guideline 5a-5c may be applied to PRM data instead. Guidelines 8 and 9 also apply for SRMs.
7. **If using DIA MS, then, if the data are analyzed with XICs, apply the above SRM guidelines (6); if the data are analyzed by extracting deconvoluted spectra, apply the above DDA guidelines 5a-5c.** DIA-MS workflows such as SWATH-MS or the equivalent on other instrument types typically yield highly multiplexed spectra that make confident identification of peptides challenging. The guidelines that apply depend on the data analysis strategy. If the data are analyzed via extracted ion chromatograms (XICs) such as with DIA-NN, OpenSWATH, Spectronaut, PeakView, etc. then the SRM guideline 6 above applies. If the data are analyzed via extracted deconvoluted spectra such as with DIA-Umpire or DISCO, then the DDA Guideline 5a-5c above applies. In addition to the raw data, the extracted deconvoluted spectra must also be submitted to ProteomeXchange repository.
8. **Even when very high confidence peptide identifications are demonstrated, consider alternate mappings of each peptide to canonical proteins other than the claimed one. Consider isobaric sequence/mass modification variants, all known SAAVs, and unreported SAAVs.** Even when a peptide identification is shown to be very highly confident, care should be taken when mapping it to a protein or novel coding element. Consider whether I=L, N[Deamidated]=D, Q[Deamidated]=E, GG=N, Q≈K, F≈M[Oxidation], or other isobaric or near isobaric substitutions, or amino acid order inversions could change the mapping of the peptide from an extraordinary result to a mapping to a commonly-observed protein. Consider if a known single amino-acid variation (SAAV) in neXtProt could turn an extraordinary result into an ordinary result. Consider if a single amino-acid change, not yet annotated in a well-known source, could turn an extraordinary result into a questionable result. Check more than one reference proteome (e.g., RefSeq may have entries that UniProtKB and Ensembl do not, and vice versa). A tool to assist with this analysis is available at PeptideAtlas at http://peptideatlas.org/map (ProteoMapper).
9. **Support Tier 1 (1A digest, 1B HLA) claims via two or more distinct uniquely-mapping, non-nested peptide sequences of length ≥9 amino acids with the above evidence in the same paper. When 2 peptides overlap, the total extent must be ≥18 amino acids. 100% coverage, except for an initiating methione, may be considered Tier 1 evidence. Tier 2A (digest) or Tier 2B (HLA) evidence may constitute just a single uniquely-mapping peptide of length ≥9 amino acids. Tier 3 evidence (not meeting Tier 2A or 2B evidence) may be discussed but is generally not considered sufficient for a claim of detection.** Single-peptide detections have too high a chance of being some type of pernicious false positive to be sufficient for claiming a new protein detection at the highest confidence. Likewise, short peptides of length 8 or smaller have relatively few peaks and have an increased chance of mapping to immunoglobulins or other variable sequences not readily apparent in the reference proteome. Nested peptides (where one sequence is fully subsumed within another) do provide additional confidence that the peptide identification is correct, but provide no additional evidence that the peptide-to-protein mapping is unique. However, microproteins can be very short, and optimal evidence can be challenging to achieve. Therefore, a tiered system of detection claims is provided. Tier 1 claims require at least 2 uniquely mapping peptides of 9 residues or longer that together cover an extent of at least 18 amino acids. For very short microproteins, 100% coverage (with the possible exclusion of the initiating methionine) can be considered Tier 1 evidence. When Tier 1 evidence is achieved via digest data (e.g. trypsin or other proteases), it is considered Tier 1A. When Tier 1 evidence is achieve in whole or in part via immunopeptidome enrichment data, it is considered Tier 1B. Lesser evidence that includes as least 1 uniquely mapping peptide of 9 residues or longer is considered Tier 2A (digest) or Tier 2B (immunopeptidome). Evidence that does not meet the above criteria may be termed Tier 3, but is generally not considered sufficient for a claim of detection. Nonethless, such “candidate detections” may be discussed to enable capture of this information by other researchers for follow up by further experiments.

---

**Supplementary Figure 1:**
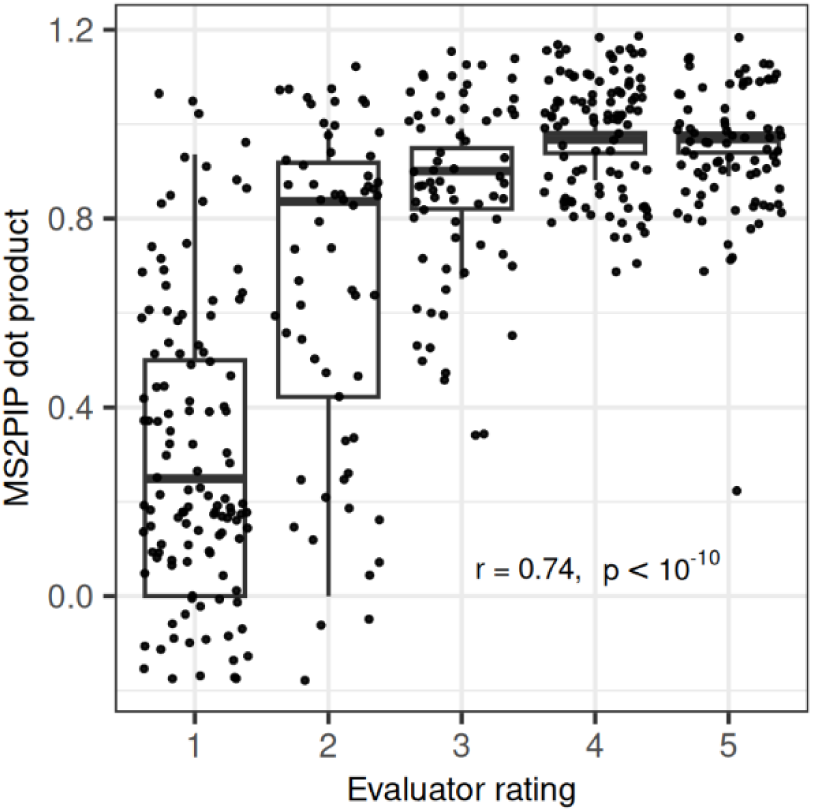
Evaluator rating is strongly correlated with dot product between observed spectra and spectra predicted by MS2PIP. MS2PIP was used to generate predicted spectra for each evaluated PSM. The dot product between the predicted and observed spectra is shown for each PSM, with PSMs grouped by manual evaluator rating. The correlation between dot products and ratings is given.

**Supplementary Figure 2:**
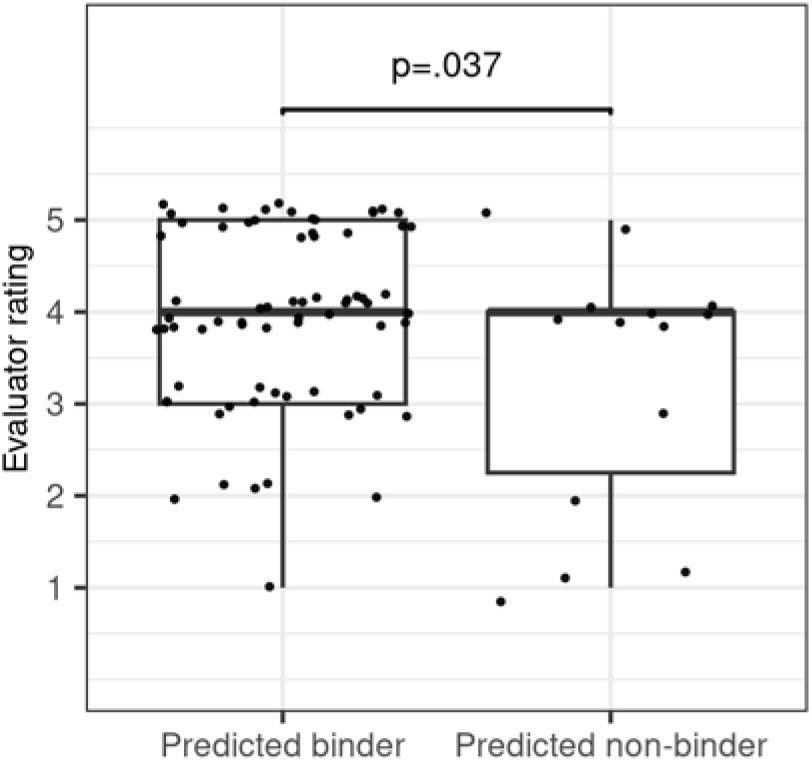
Peptides that are predicted to bind HLAs are more highly rated by evaluators. For each evaluated immunopeptidomics peptide from Ouspenskaia et al. 2021, Martinez et al. 2020, or Chong et al. 2020, HLA binding was predicted using NetMHC. Any peptide meeting the NetMHC criteria for weak or strong binder to an HLA allele present in the cell type was considered a predicted binder. The distribution of evaluator ratings among predicted binders and peptides not predicted to bind HLAs is shown.

**Supplementary Figure 3:**
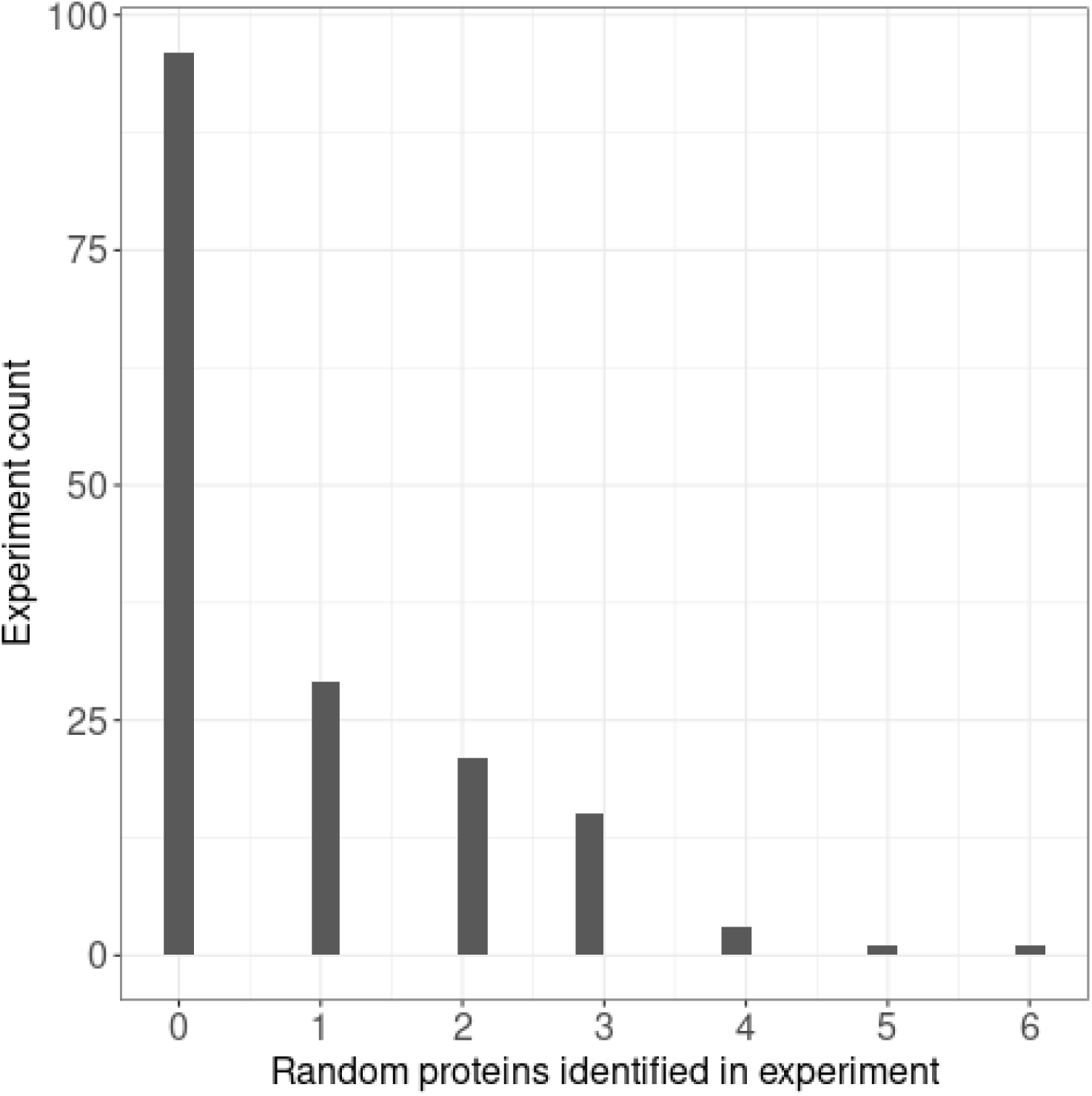
Proteomic searches for random proteins in human MS datasets falsely report detections when canonical proteins are excluded from the protein database. Histogram showing the number of random proteins detected among studies when MSGF+ was used to detect a sample of 10 randomly constructed proteins against a human MS dataset with 166 experiments. Proteins were considered detected if they had a peptide with a reported q-value <1%. This plot demonstrates that, in the absence of genuine detection of any protein in the database, it is common for a few proteins to be reported with q-value <1%. This is because the q-value for a given PSM is estimated as the number of decoys with confidence score above that of the PSM divided by the number of targets with confidence scores above the PSM. Under the null hypothesis of zero genuine detections, it is equally likely that a target or decoy has the highest confidence score; when it is a target it will be assigned a q-value of 0. For instance, Chothani et al.^4^ employed a two-stage strategy to detect sORF products. In the first stage, the UniProt human proteome was used as the sequence database. For each MS experiment, any spectra that matched with a peptide at the 1% FDR threshold was removed from the spectra file. In the second stage, the sORF list was used as the sequence database against the modified spectra file, and any sORF product with a peptide identified at the 1% FDR threshold was considered to be detected. Since all annotated proteins were removed from the database in the second stage, and there may be no unannotated proteins detectable in the sample, the conditions of no genuine protein detections are potentially met. As shown by this plot, under these conditions false positives are expected.

**Supplementary Figure 4:**
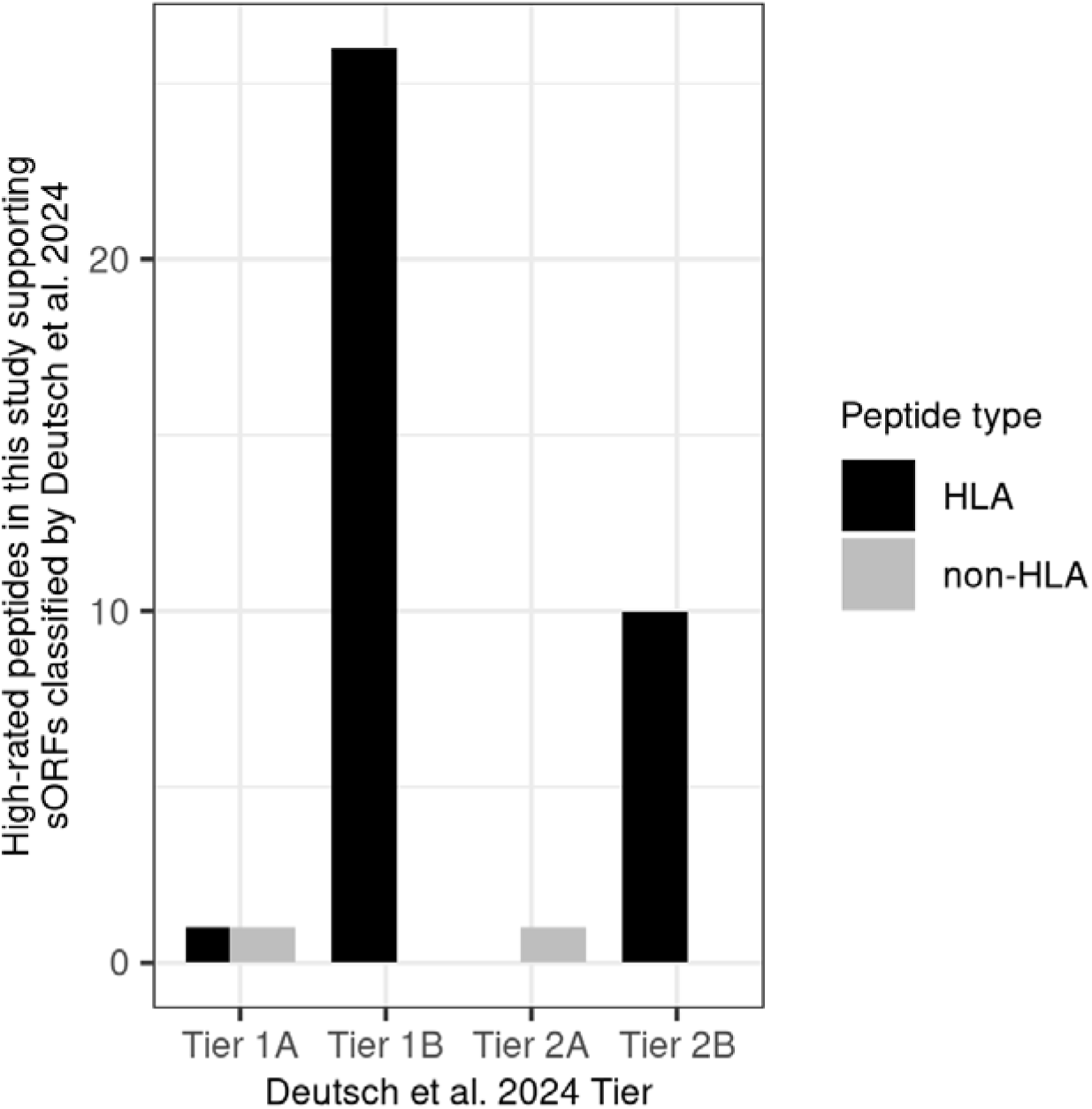
Peptides highly rated in this study that also supported ORFs classified in Deutsch et al. 2024. The number of high-rated peptides (ratings of 4 or 5) analyzed in this study that were also validated in Deutsch et al. 2024 and used as support for ORFs of various tiers classified in that paper. Peptides are divided by whether they were found in HLA immunopeptidomics experiments or non-HLA conventional proteomics experiments. The ORF tier definitions, taken from Deutsch et al. 2024, are as follows. Tier 1A: ORF supported by at least two non-nested peptides from conventional proteomics experiments as well as Ribo-Seq data. Tier 1B: ORF supported by at least two non-nested peptides from HLA immunopeptidomics experiments as well as Ribo-Seq data. Tier 2A: ORF supported by at least one peptide from conventional proteomics experiments as well as Ribo-Seq data. Tier 2B: ORF supported by at least one peptide from HLA immunopeptidomics experiments as well as Ribo-Seq data.

